# A scalable culturing system for the marine annelid *Platynereis dumerilii*

**DOI:** 10.1101/714238

**Authors:** Emily Kuehn, Alexander W. Stockinger, Jerome Girard, Florian Raible, B. Duygu Özpolat

## Abstract

*Platynereis dumerilii* is a marine segmented worm (annelid) with externally fertilized embryos and it can be cultured for the full life cycle in the laboratory. The accessibility of embryos and larvae combined with the breadth of the established molecular and functional techniques has made *P. dumerilii* an attractive model for studying development, cell lineages, cell type evolution, reproduction, regeneration, the nervous system, and behavior. Traditionally, these worms have been kept in rooms dedicated for their culture. This allows for the regulation of temperature and light cycles, which is critical to synchronizing sexual maturation. However, regulating the conditions of a whole room present limitations, especially if experiments require being able to change culturing conditions. Here we present scalable and flexible culture methods that provide ability to control the environmental conditions, and have a multi-purpose culture space. We provide a closed setup shelving design with proper light conditions necessary for *P. dumerilii* to mature. We also implemented a standardized method of feeding *P. dumerilii* cultures with powdered spirulina which relieves the ambiguity associated with using frozen spinach, and helps standardize nutrition conditions across experiments and across different labs. By using these methods, we were able to raise mature *P. dumerilii*, capable of spawning and producing viable embryos for experimentation and replenishing culture populations. These methods will allow for the further accessibility of *P. dumerilii* as a model system, and they can be adapted for other aquatic organisms.

## BACKGROUND

Nereidid worms such as *Platynereis* have been popular in studies of development and fertilization because of transparent, abundant, and comparatively large eggs and embryos [1,2]. As researchers like Edmund Beecher Wilson did in the late 19th century, many labs today benefit from *Platynereis* as a model organism for addressing a wide range of biological questions such as cell type evolution, nervous system evo-devo and activity, reproductive periodicity, circalunar cycling, endocrinology, regeneration, post-embryonic segment addition, stem cell biology, fertilization, oocyte maturation, embryonic and larval development [3–16]. The sexual worms broadcast spawn, producing thousands of externally-developing embryos. The embryos are large enough to inject but small (about 160 µm in diameter) and transparent enough to image live and fixed samples. *Platynereis* has a relatively quick and highly synchronized embryogenesis: it takes only 18 hours from fertilization to hatching as a planktonic trochophore larva in *P. dumerilii [2,17]*. This allows researchers to study embryonic development over the course of just one day. A number of tools and techniques are already established in *P. dumerilii* [3] including microinjection [5,12,18], transgenesis and genetic tools [18–21], single cell RNA sequencing [22], behavioral tracking [8,16], and live imaging [5]. This well-equipped tool kit combined with the large number of embryos generated by each fertilization make *P. dumerilii* an attractive model organism which can be cultured under specific laboratory conditions for its full life cycle.

Ernest E. Just was among one of the earliest of people who tried rearing *Platynereis* in the lab at the beginning of the 20^th^ century [23]. He studied fertilization in *Platynereis megalops*, which he collected from Great Harbor in Woods Hole [24], and wanted to have access to eggs throughout the year instead of only during the summer. In Europe, studies of *P. dumerilii* go back to early 20^th^ century [25] at Naples Zoological Station. *P. dumerilii* cultures today are thought to mostly originate from the Bay of Naples, and have been bred in the lab since 1950s (originally by Carl Hauenschild) [2,17]. Even though resources for culturing *P. dumerilii* exist including Fischer and Dorresteijn’s excellent guide online [26], these resources only provide guidelines for larger scale culturing of *Platynereis* and detailed guidelines for establishing small (but scalable) culturing are not available. For many research areas (such as physiology, behavior, aging, reproduction…) there is also the need for flexibly adjusting the environmental parameters, which is challenging with the traditional methods of culturing *P. dumerilii*. Finally, several areas for standardizing culturing methods remain to be established, especially regarding the feeding methods. This is particularly important for studying biological processes are affected dramatically by nutrition, and being able to carry out comparable experiments across different research labs.

Here we describe a scalable, small footprint setup for culturing *P. dumerilii*, including detailed methods of light regulation, light and temperature monitoring, husbandry, and feeding. This setup, with blackout curtains and its own automatic lighting which serves as the sun and moon, removes reliance on dedicated culture rooms. It provides greater flexibility for choosing and adjusting culturing components (such as light source) and can be put together at a lower cost than similar designs that use incubators in place of a shelving unit [27]. We provide worm maturation data that can be used to scale the culture up or down, based on the number of mature worms needed. We also present standardized feeding methods we developed using powdered, commercially available nutrients such as spirulina. This homogenous suspension can be distributed throughout the culture boxes evenly. Our feeding method eliminates the ambiguity associated with using fresh or frozen spinach, and will enable better comparison of data across laboratories especially for types of research, such as physiology, where diet is a particularly important factor. Finally, we present data and images of *P. dumerilii* embryos, larvae, juveniles, and adults we obtained from our cultures, and visualization of normal development, confirming the robustness of the culturing conditions.

## METHODS

### Water Type and Filtering

*P. dumerilii* embryos, larvae, juveniles and adults were kept in full strength natural filtered sea water (NFSW). The sea water was first filtered through a 1 µm filter system in the Marine Resource Center at the Marine Biological Laboratory. The 1 µm NFSW was used primarily for juvenile and adult culture boxes (Suppl. Fig. 1A).

For food preparation, microinjections, and culturing embryos and young larvae, sea water was filtered further via a glass graduated filtration funnel (*Pyrex*, 33971-1L) secured to a glass funnel stem (*Sigma*, Z290688) by a spring clamp (*Millipore*, xx1004703) (Suppl. Fig. 1). On top of the funnel stem, a 0.22 µm pore size nitrocellulose membrane (*Millipore*, GSTF04700) was placed, and this membrane was covered by one piece of Whatman filter paper (*GE Healthcare*, 1002055). The filter unit was placed on a glass vacuum flask (*Fisherbrand*, FB-300-4000) fitted with a rubber adapter (*Fisherbrand*, 05-888-107) so that the stem sat snugly in place (Suppl. Fig. 1B).

**Figure 1.**
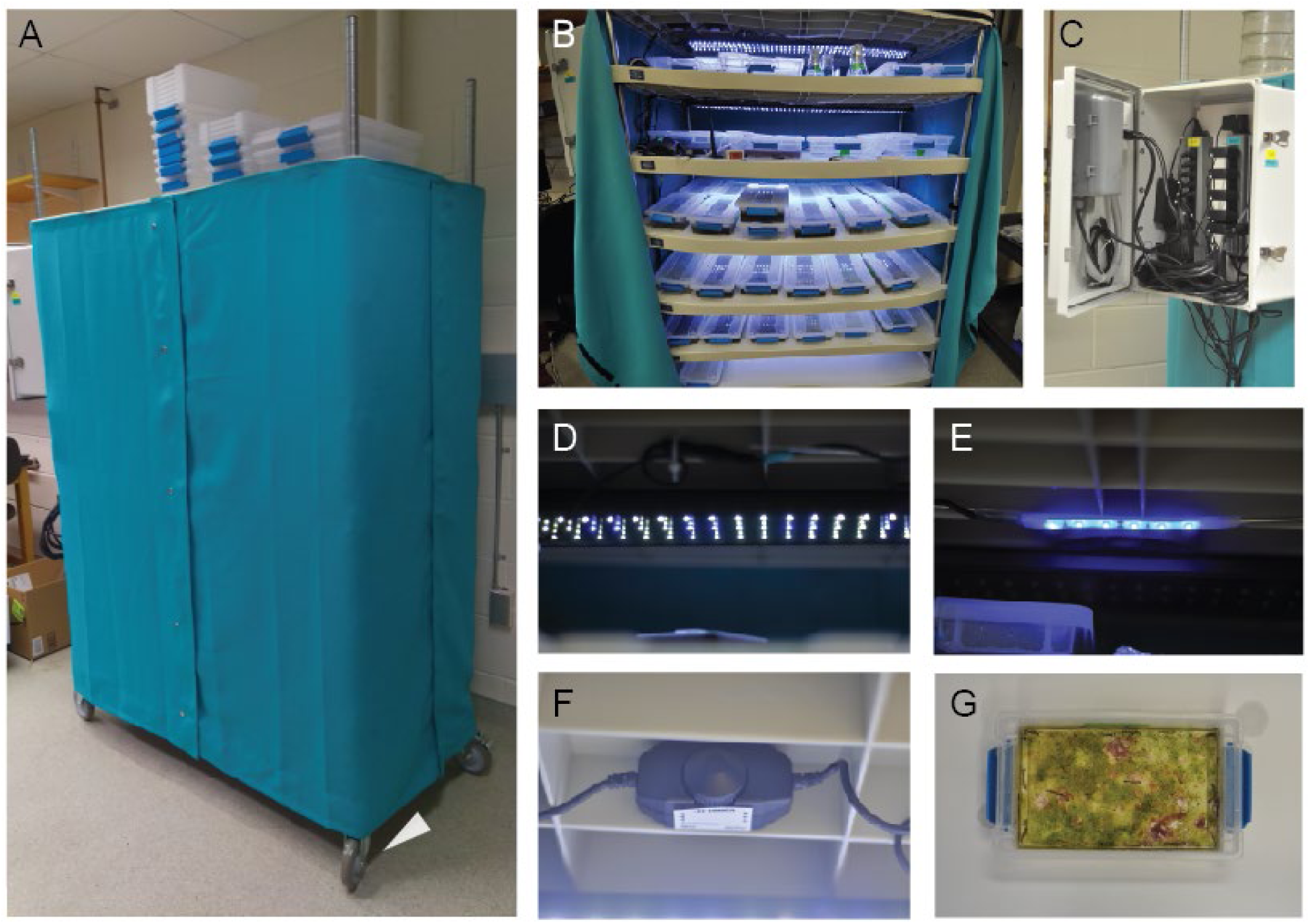
Scalable culture setup for *Platynereis dumerilii*. A) The shelving unit is enclosed in a blackout curtain which keeps unwanted light out during “moon off” periods of *P. dumerilii*’s lunar maturation cycle. Putting the unit on wheels (arrowhead) allows us to easily access culture boxes at the back of the shelves. The curtain can also be opened from both sides. B) Each shelf within the unit is equipped with identical lighting to ensure all culture boxes are receiving both sun and moonlight. C) The lighting is controlled automatically. Controls are located in a box outside of the unit. D) Sun lighting on. E) Moon lighting on. F) The intensity of moonlight can be easily adjusted with the dimmer shown here. A wireless light monitor was used to ensure each shelf received equal light intensity. G) Sterilite boxes made of polypropylene and polyethylene were used to house worms. After several months of culturing, algae was seen to grow in the boxes without harmful effects to the worms.

### Collecting embryos for sustaining cultures

To spawn mature worms, male and female worms were placed together in a small glass dish of approximately 150 mL 0.22 µm NFSW. After spawning, worms were removed from the dish, sea water with excess sperm in the dish was discarded and replaced with clean sea water to prevent polyspermy, and embryos were kept at 18°C. Around 24 hpf, larvae were poured onto a 85 µm sieve and 0.22 µm NFSW was poured over the embryos several times to wash the jelly off. To make a 85 µm sieve, we cut off the conical end of a 50 mL tube (and removed the cap), stretched a 85 µm pore mesh (*Component Supply*, U-CMN-85-A) over one of the ends of the tube, and secured the mesh with a rubber band or glue.

After the jelly was washed off, larvae were pipetted off the sieve and into a clean dish of 0.22 µm NFSW, or the sieve was flipped over an empty dish and larvae were washed into the dish. Penicillin Streptomycin (*Gibco*, 15140-122) (1:1000) was added, and the cultures were kept at 18°C until 7-10 days post fertilization (dpf).

### Setting Up Culture Boxes – High and Low Density Populations

As containers for culturing, Sterilite brand boxes made of polypropylene and polyethylene (PP5 type non-reactive plastic) were used (Fig. 1G). These boxes are non-toxic, inexpensive, come in several sizes, and have lids that can sit loosely to allow air in and out when the latches are not used. To ensure no residual chemicals or dust from production remained, the boxes were rinsed with deionized water, soaked overnight, and washed with a clean sponge without any detergent before they were used for cultures for the first time.

Healthy larvae at 7-10 dpf were transferred into Large Sterilite boxes (*Sterilite*, 1963) (no air bubbler needed at this stage), and were kept in the 18°C incubator until around 1-2 month(s) post-fertilization (around the time young juvenile worms started building tubes). At this time, these cultures were transferred from the incubator to the shelving unit to begin assimilation to the lighting schedule. These large boxes contained a high-density population of worms of about 300-350 per box. To facilitate the dispersal of oxygen throughout these cultures with growing worms, high density boxes were equipped with air bubblers (*Tetra*, Whisper Air Pump, 77846) once transferred to the shelf. Air tubes were inserted by making a hole on the lid using a hot glue gun tip (using the heat to melt the plastic, without the glue).

Low density cultures were established a little over two months (typically at the beginning of the third month) after the worms were born. Using a paint brush, the worms were pushed gently to come out of the tubes and were collected with a pipette to be transferred to a low density box. These cultures consisted of 30 worms per small Sterilite box (*Sterilite*, 1961) in 500 mL 1-µm-filtered natural seawater, and were not aerated.

### Feeding *Platynereis* larvae

Depending on availability at our facilities, we used either *T-iso*, or *T. chuii*, or a mix of both algae species to feed the young *P. dumerilii* larvae (starting around 7-10 dpf). The algae cultures were kept in a room with ample natural light and/or with additional LED lights. Algae cultures were grown in glass or plastic carboys and were aerated to ensure faster growth. Algae were collected into 50 mL centrifuge tubes when cultures were seen to be dense enough (dark brown or green color, depending on the species). The tubes were then centrifuged for 10 minutes at 2000 rpm. The supernatant was discarded and the tube was refilled with additional algae stock. Centrifugation was repeated until the pellet is large enough to fill the conical part of the tube (∼2.5mL). Then algae pellet was resuspended in 50 mL of 0.22 µm NFSW. This provided concentrated algae stocks which could then be stored at 4°C for later use. During the first month of development, 25 mL per large box (1.5 L water volume) of this concentrated stock was distributed using a transfer pipette, twice per week. On the second month of development, the juvenile worms were switched to the spirulina regimen (see Results and Discussion below for details).

### Maintaining the larval and juvenile culture boxes

#### Maintaining larvae

The larvae may perish easily if water cleanliness is not maintained. To ensure optimal water cleanliness larval culture boxes were checked under the microscope regularly. If growth of protozoans was observed, water was removed using a sink-enabled vacuum through a 85-µm sieve which is large enough to let protozoa out but prevents removing small larvae. 0.22µm NFSW was used to culture the larvae.

#### Maintaining juveniles

The NFSW in the culture boxes was completely replaced every two weeks with new 1.0 µm NFSW. For boxes with juvenile worms that have already formed tubes, dirty water was poured completely into a plastic dishpan, as the worms mostly stay in tubes. Any escaped worms were transferred back into the culture box from dishpan. The dirty water was discarded into a container to which bleach was added prior to final disposal into the drain, in order to prevent introducing *P. dumerilii* into the environment. For cultures with very young juveniles, dirty water is removed using a vacuum filter (as with larval cultures above).

### Mature worm collection and maintenance

Mature worms were collected and separated into females and males in large Sterilite boxes equipped with air bubblers. Mature worms for which the gender could not be identified (“unknown gender” worms) were placed with the males, as males are thought to not spawn in the presence of immature females. The boxes were monitored daily to remove dead worms, change the water if needed, and remove and use mature worms for setting up new cultures and for experiments. Mature worm collection was done systematically only during 2 weeks (on Mondays, Wednesdays, and Fridays) in a month when the worm maturation is expected to peak (starting about 10 days following the last day of “moon on”).

### Temperature Control and Monitoring

Most labs use 18°C as the culturing temperature for *P. dumerilii* for the full life cycle. Previous studies on embryonic and larval development suggest that at least for these early stages the animals can be kept at lower or higher temperatures (14-30°C) [17], while a systematic testing of temperatures higher than 18°C for the full life cycle has not been reported. We have found that our cultures tolerated slightly higher average temperatures (19-20.5°C) (Suppl. Fig. 2). The thermostat of the culture room was set to 20°C. However, we kept an additional portable AC unit (*Arctic King*, WPPH08CR8N) set to about 18°C (65°F) running next to the cultures, which served as a back-up. We also used a large 18°C incubator (no light cycles) for keeping some worm cultures as reserves, in case the room temperature control severely failed.

To monitor temperature, a Monnit wireless temperature monitor (*Monnit*, MNS-9-IN-TS-ST) was placed on one of the shelves. In addition, we also used a thermometer (*Suplong*, COMINHKPR144821) which stores the minimum and maximum temperatures recorded within the shelving unit for manually checking the temperature fluctuations. The data collected by the Monnit thermometer was stored online and provided a complete list of all temperature readings. This system also sends alerts if it detects temperatures outside of a specified range.

### Light Spectra Measurements

For the sun (*Nicrew*, ZJL-40A) and moon (Ebay seller: *21ledusa*, 700381560185) light sources, the irradiance per wavelength was measured using a spectrometer (*International Light Technologies*, ILT-ILT950-UV-NIR). To achieve different light intensities, an LED dimmer (DC12V∼24V, *Supernight*) was used. The spectrometer was set up at 20 cm distance to the light source in an otherwise dark room. For plotting the data, the sensor output (in µW/cm^2^) was converted to the more commonly used photon flux (photons/(m^2^*s)).

## RESULTS AND DISCUSSION

### Building the scalable shelving unit

The first consideration for a lab to begin culturing *P. dumerilii* is the availability of appropriate housing infrastructure. In nature, synchronized reproduction occurs in phase with the lunar cycle. To mimic this, *P. dumerilii* is typically maintained in a defined light regime, consisting of 16 : 8 hours of daylight : night, with dim nocturnal lighting simulating a full moon stimulus for several nights within a month (see below). Most labs currently working with *P. dumerilii* keep their worms in a separate culture room and control the room lights to achieve day, night, and moon conditions. But having a dedicated worm culture room may not always be feasible or even needed if only a small scale of *P. dumerilii* culturing is desired. Additionally, *P. dumerilii* maturation peaks for only two weeks out of the month due to the lunar maturation cycle. Labs wanting to have mature worms continuously would need separate culture spaces on opposite lighting schedules.

To circumvent the need for separate culture rooms, and to use the space available for multiple purposes, we designed a stand-alone culture setup (Fig. 1) (Suppl. File 1 for a comprehensive list of all the parts, reagents, and ordering information). The setup is composed of a shelving unit (*Nexel*, 188127) with wheels (*Nexel*, CA5SB) for ease of access, black-out curtains to prevent unwanted light reaching the worms during the night hours when complete darkness is needed (i.e. no “moon”), and two sets of lights (sun and moon) on separate circuits installed on each shelf controlled by a timer (Fig. 1C, Suppl. File 2). With this setup, labs wanting to have mature worms available during the entire month could have two or more culture units on opposite moonlight schedules in the same room.

The shelving unit (Fig. 1A, B) was assembled according to the manufacturer’s directions. The unit comes with 4 shelves. We added 3 more shelves (*Nexel*, S2448SP), each being approximately 8 inches (20.3 cm) apart from each other. This allowed enough space for the lights and stacking up 2 rows of culture boxes. Small (low worm density) boxes can fit into two rows on each shelf of our culturing unit. They can also be stacked, thus in theory 28 small Sterilite boxes (14 if unstacked) can be stored per shelf.

After assembling the shelving unit, the sun (*NICREW*, B06XYKD67V) (Fig. 1D) and moon (Ebay seller: *21ledusa*, 700381560185) (Fig. 1E) lights were installed on each shelf by punching holes on the plastic and using aluminum welding rods to secure the lights (Figs. 2E-E’, 3B). See Figures 2 and 3 for circuit diagrams and blueprints for the assembly of lights. Dimmers (*HitLights*, B00RBXPDQU) were included in the circuits to control the brightness from each set of lights (Fig. 1F, 2B).

**Figure 2.**
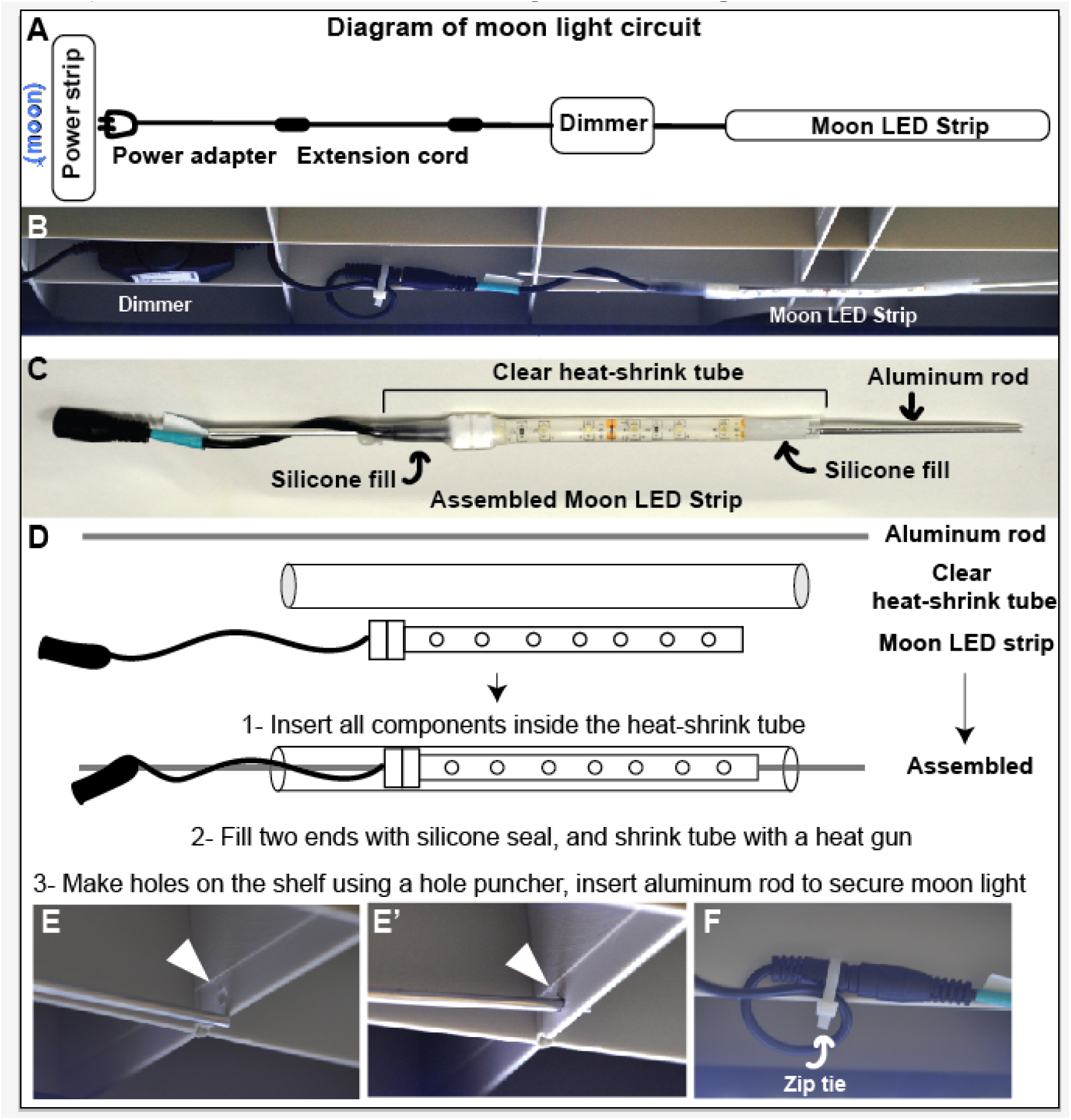
Setting up the moon light. A) Diagram of moon light circuit. One moon light per shelf is installed, connected to a dimmer and an extension cord (for reaching to the power strip). The moon power adapters should be plugged into the power strip controlled by the timer channel dedicated to the moon lights. (Note that power adapters come with the LED strips and do not need to be purchased separately.) B) Photograph showing the dimmer and LED strip components, attached to the shelf. C) Components of the moon light LED strip. The LED strips are purchased and then modified for the shelving. Aluminum rod helps with attaching the light to the shelf. Clear heat shrink tube and silicone is for sealing the light for protection from water. D) Schematic showing the components of the moon LED light, and assembly instructions. E) Using a hole puncher, plastic shelving is modified for insertion of the aluminum rod. Arrowheads: holes F) Holes can also be used for securing cables with zip-ties.

**Figure 3.**
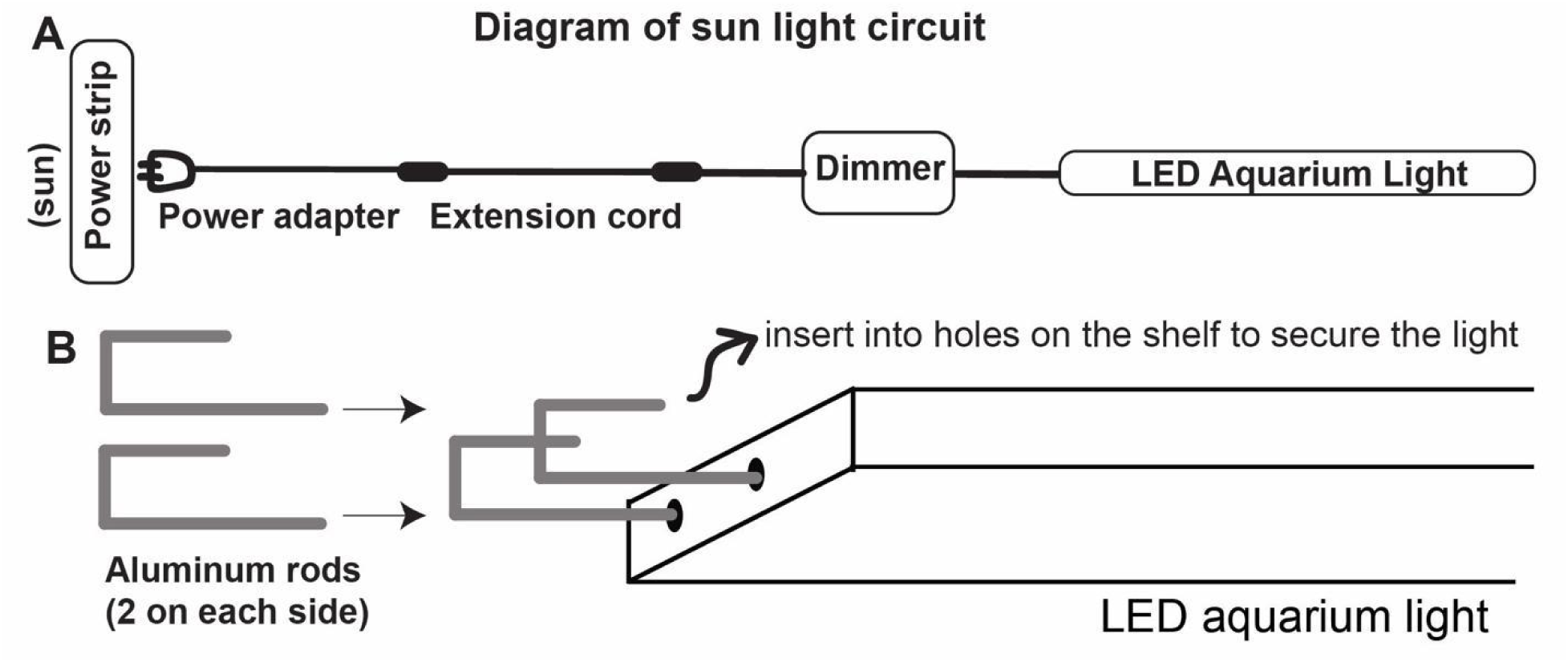
Setting up the sun light. A) Diagram of the sun light circuit. One aquarium light per shelf is installed to provide light during the day. The aquarium light is connected to a dimmer and an extension cord (for reaching to the power strip). The power adapters should be plugged into the power strip controlled by the timer channel dedicated to the sun lights. In addition, aquarium light switch should always be ON, the turn ON-OFF will be controlled automatically by the timer. B) Bent aluminum rods (2 on each side) are used for securing the light onto the shelf. See Figure 2E-E’ for aluminum rod insertion and hole punching.

Next, 4 pieces of blackout curtains (*Deconovo*, B01MU1CMSD, 42×63 inches) were modified by sewing hook and loop (e.g. Velcro) strips on the curtains along the edges (Fig. 4). We found that using 4 pieces of fabric provided flexibility of access to cultures from all sides of the shelving unit, as well as ease of running cables from the lights and air pumps to the power box (Fig. 4B). A basic sewing machine (Singer Simple 3232) and needles made for thick fabric were used for sewing the hook and loop strips onto the curtains.

**Figure 4.**
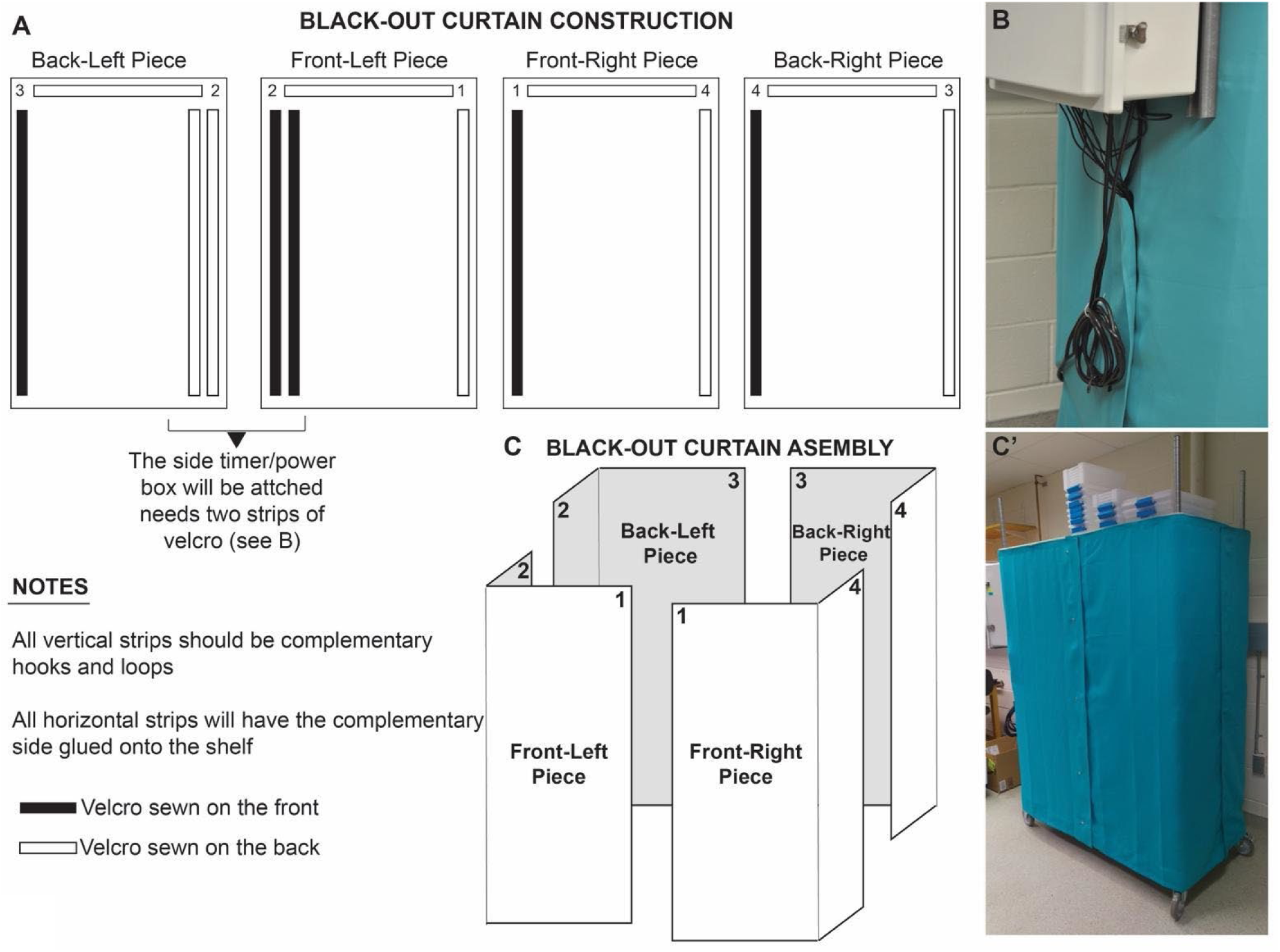
Blackout curtain construction and assembly. A) Instructions for sewing Velcro (hook and loop) strips onto the blackout curtains. Top parts of the curtains will have one side of adhesive Velcro sewn on, and the other side applied onto the shelf. Even if adhesive type of Velcro is used, the adhesive alone is not sufficient to keep Velcro strips on fabric, and these strips need to be sewn. B) The side of shelving that will have the power box installed will have two Velcro strips to allow for extra sealing for cables from the shelf to run outside to the box. C-C’) Schematic showing the assembly of 4 curtain pieces to cover the shelving unit (C) and a picture of the assembled shelf (C’). Note that the uppermost shelf is used for storing culture boxes, and does not function for culturing worms as it is not covered with curtains.

After the blackout curtains were installed, the power box (*Allied Moulded Products Inc*. AM2068RT) was attached to one of the shorter sides of the shelving (either left or right side would be fine, depending on which side is more comfortable for a given space). The power supply box was attached to the shelves using two plastic struts (on the left and the right of the back panel) sandwiched between the box and shelves; the struts were helpful to support the screws securing the box onto the shelf. Next, power strips and the timer were installed into the power box (Fig. 1C). The sun and moon lights were connected to the timer in separate circuits with the aid of an electrician.

### Light/Dark Cycles

*P. dumerilii* mature according to a lunar cycle. In their natural habitat, worms will congregate at the surface of the water within about ten days following the full moon for a period of about 2 weeks to spawn [25,28]. In the lab, a synchronized swarming pattern can be obtained by simulating 28-day lunar cycle lighting conditions [16]. This is achieved by having a period of 8 nights where a dim light turns on at night in order to mimic the moon. This period is followed by 20 days in which the worms are kept in complete darkness during the night hours. Note that the “moon” lights do not mimic the phases of the moon; worms respond to simple presence/absence of light at night. Setting the lunar lighting to a schedule of four exact weeks simplifies planning in terms of knowing when the worms will peak in maturation. We followed these standards established by others before in our culture conditions: For all days, worm cultures in the shelving unit received 16 hours of daylight and 8 hours of night (either with the moonlight on or off) (Fig. 5A). These on and off cycles were controlled automatically by an industrial timer (*MRO Supply*, DG280A), and settings were adjusted according to manufacturer’s user guide (Suppl. File 2 for timer settings). The timer removed the need for manually installing and uninstalling the moon lights.

**Figure 5.**
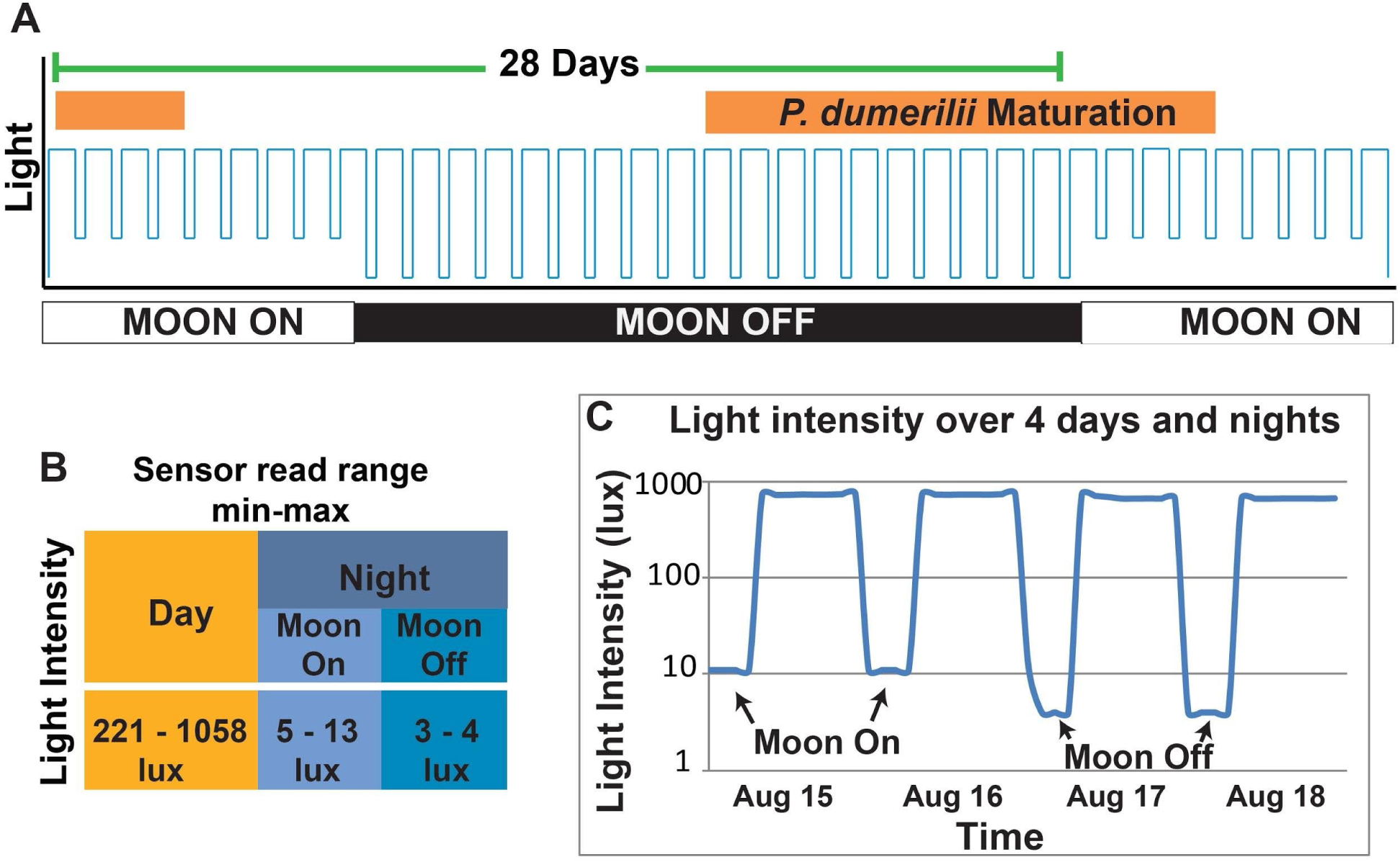
Lighting Schedule. A) The schematic shows the lighting schedule we used to culture *P. dumerilii* through maturity. Worms received 16 hours of sunlight every day and 8 hours of either moonlight or complete darkness at night. In a 28 day cycle, the moon is on for 8 nights and off for the remaining 20 days. *P. dumerilii’s* maturation period (indicated with the orange bar) begins 10 days after the last night the moon was on and lasts for 2 weeks. B) Values of light intensity (lux) were collected by a wireless light sensor. The sensor was placed inside a culture box in order to record the amount of light the worms were receiving through the translucent culture box lids. C) Example light sensor data recording over 4 days showing the transition between a period when the moonlight was on and off.

To monitor that light conditions were properly set and maintained during the day and night, we used a Monnit wireless light monitor (*Monnit*, MNS-9-W2-LS-LM). The monitor sends the data to a wireless receiver installed at our facility, and users can access the sensor reads online via a simple user interface (https://www.imonnit.com/). The light monitor was periodically moved between shelves to ensure that the light signal reaching the worms was the same throughout the unit and that all the sun and moon lights were functioning properly. We then continued to regularly monitor each shelf lighting this way for any lights that may need replacement.

Using the light monitor, we collected several types of light intensity data. To measure the amount of light each shelf received, we rotated the light sensor between shelves recording light levels for 24 hours each rotation (Fig. 5B-C). This was done for a period of two months so that we could have several readings from each shelf during “moon on” and “moon off” periods. For the first month, the sensor was placed on top of a Sterilite box. For the following month, the monitor was placed inside an unused small Sterilite box to get a more accurate estimate of the amount of light the worms received through the translucent lids. Of the measurements taken from inside a Sterilite box, the daylight range was between 221-1058 lux (see below for detailed photon flux information). Even though we did not systematically test whether the lowest or highest settings of illumination had different effects, we obtained mature worms from all shelves. Thus we conclude that values within this range will be sufficient for the worms to grow and mature.

The moonlight range measured from inside a Sterilite box was 5-13 lux during a “moon on” period and 3-4 lux during a “moon off” period (Fig. 5B). To determine whether the reading of 3-4 lux during a moon-off period was due to sensor error or an outside light which infiltrated the unit’s curtains, we placed the sensor in a completely sealed and opaque box, and the sensor still read 3-4 lux. Therefore, we assume that when the sensor read 3-4 lux, there was no or negligible light within the culture setup. Taking these values as “zero” for this particular sensor, we adjusted the moonlight brightness to a higher range of brightness using the dimmers aiming for 10-15 lux (Fig. 5C), taking previously-published values into account [16].

### Light spectra distribution of the moon and sun lights

The distribution and strength (irradiance) of the wavelength available to organisms can affect many biological processes such as circadian rhythm, sexual maturation etc [29,30]. We next determined the spectral distribution of the moon and sun lights in our culture setup by using a photo spectrometer. We tested the spectral properties of both light sources at different light intensities and measured the energy flux for each wavelength. We found that the moon lamp consistently emitted light around 450 nm wavelength at different intensity settings (one example shown in Fig. 6A). We also found that the sun lamp emission spectrum did not change with at two different levels of illuminance tested: 200 lux (Fig. 6C) and 1000 lux (Fig. 6D). Overall, these results indicate that these light sources can be used at any intensity needed for a given experiment without causing a change in the emission spectrum.

**Figure 6.**
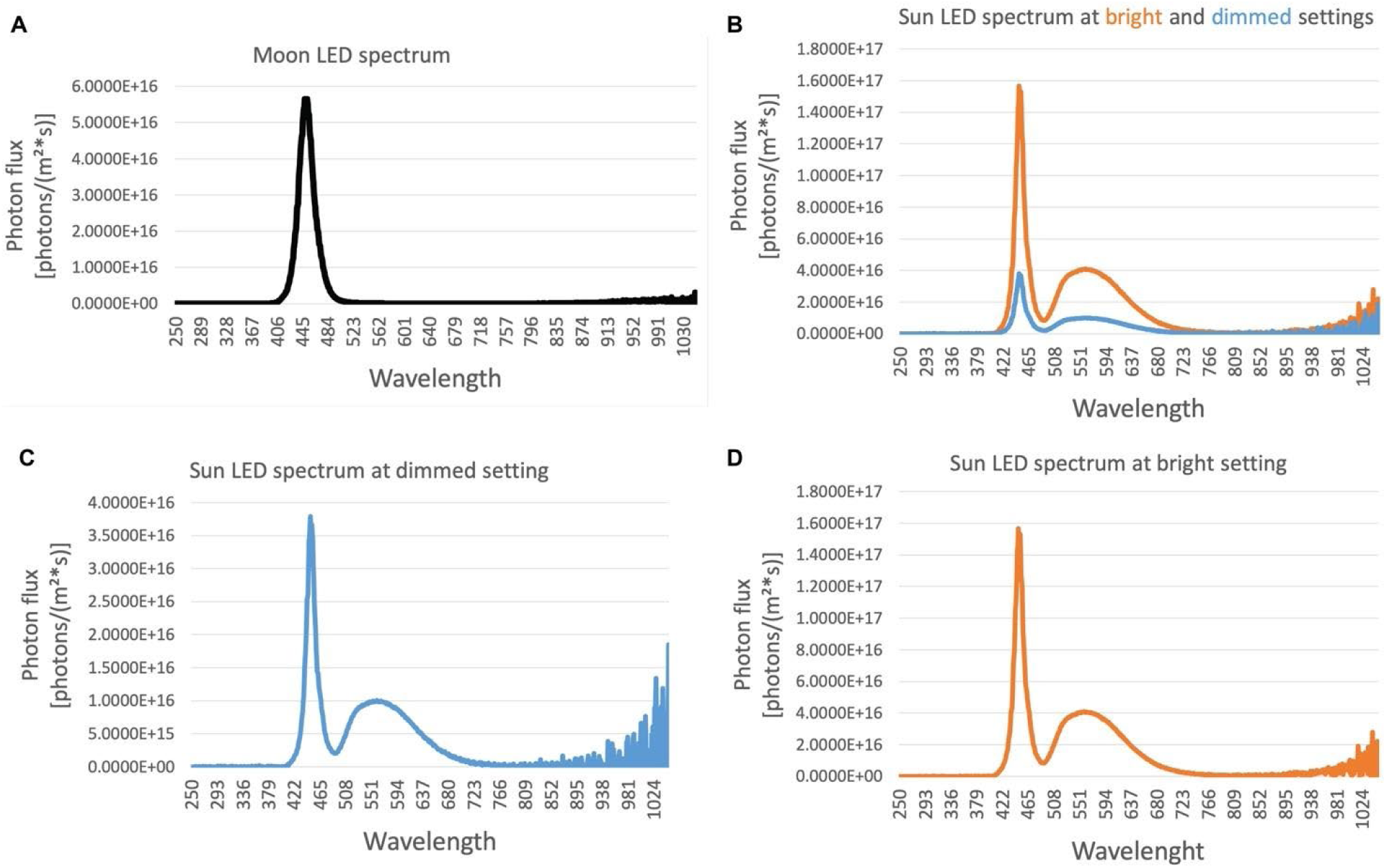
The distribution of irradiance per wavelength of the moon and sun LEDs. A) An example of moon LED light irradiance per wavelength is shown. The emission was around 450 nm at the different light intensities tested. B) Sun LED light spectral distribution did not change when the brightness of the lamp was changed between ∼1000 lux (bright) and ∼200 lux (dimmed). Graphs in (C) and (D) show the dimmed and bright setting distributions in (B) separately.

### Developing a standardized feeding method for juvenile *Platynereis dumerilii*

To date, most labs that culture *P. dumerilii* have fed them frozen spinach (organic), tetraselmis fish flakes (*Tetra*, 7101), and live phytoplankton [27]. We found traditional methods to be ambiguous in terms of how much spinach the worms were actually receiving (e.g. “5 gr frozen spinach” could contain variable amounts of actual spinach, depending on the frozen water content of the product). Also studying growth and other physiological processes require standardized feeding conditions that can be replicated across different labs.

We therefore set out to develop an easy-to-replicate method of feeding (Fig. 7): in essence, cultures were given powdered spirulina (1.0 g/L) (*Micro Ingredients*) and Sera micron flakes (0.3 g/L) (*Sera*, 0072041678) suspended in 0.22 µm NFSW (also see Suppl. File 1 for recipes and volumes of food used per box size). This way the worms received a homogenous mixture of food, the volume of which could be easily adjusted if fouling was observed or if food was consumed too quickly (Suppl. Fig. 3). Small, low density boxes (30 worms) were fed with 20 mL of this mixture and the larger, high density boxes (>100 worms) received 40 mL. Worms were fed twice per week, on Tuesdays and Fridays (note that labs using the spinach-tetramin regimen typically feed their cultures on a Monday-Wednesday-Friday schedule).

**Figure 7.**
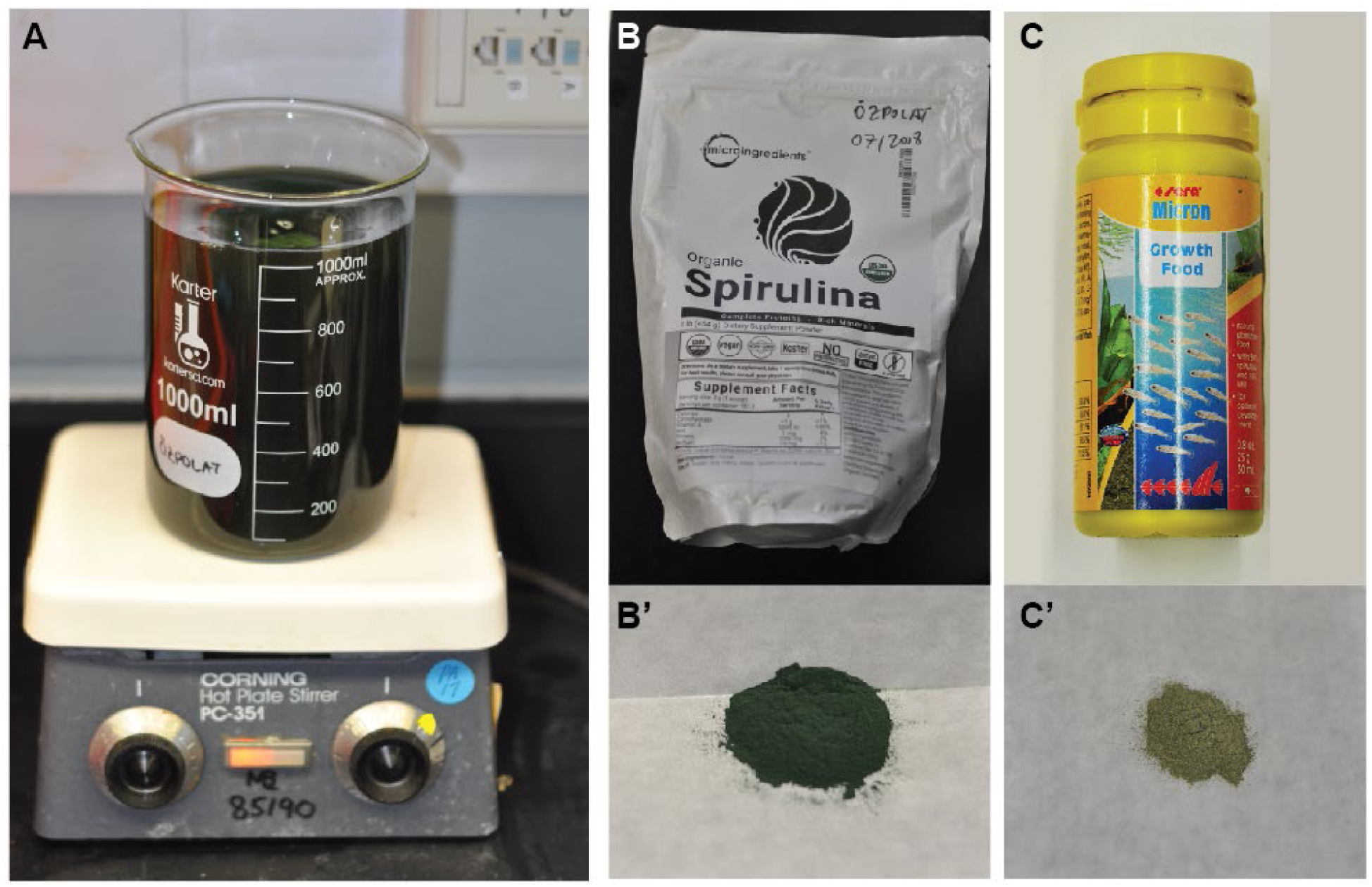
Standardized feeding methods. A) 1 g spirulina and 0.3g sera micron flakes were added to 1 L FSW and mixed thoroughly to create a homogenous solution. B-B’) Spirulina powder obtained from Microingredients. C-C’) Sera micron growth food.

### Comparison of spinach and spirulina feeding regimens in juveniles

To compare the new spirulina-sera micron cocktail feeding method to the traditional spinach-fish flakes-algae feeding regimen, we tested the growth rate difference between groups of juvenile worms that were on either of these diets. For simplicity we refer to these as spinach versus spirulina feeding regimens, even though the spinach regimen also includes fish flakes and algae, and the spirulina regimen includes sera micron. This experiment was carried out at Florian Raible’s laboratory at MFPL (Vienna), where the primary feeding regimen is the spinach regimen.

For the experiment, 80 sibling juveniles (strain PIN619512 R-mix) that were 53 days old were split into eight culture boxes (500 mL, 1:1 AFSW:NFSW). Half of the boxes were fed with spinach regimen, and the other half with spirulina regimen. The spinach-fed animals received 0.5 grams of organic spinach leaves every Tuesday, and 10 mL of algae cocktail (containing 0.25 g/L finely ground Tetramin flakes in lab-cultured *Tetraselmis marina* algae solution) every Friday. These values are based on estimated averages used by the Raible Lab, since a quantifiable feeding regimen has not been established. The spirulina-fed animals received 10 mL of spirulina cocktail every Tuesday and Friday (same spirulina regimen recipe as reported above and in Fig. 7).

At the time of setting up the experiment (t=0) worms were anesthetized in 1:1 NSFW and 7.5% Magnesium Chloride (MgCl2) [12] and the number of segments were counted for each individual (average number of 19.2 segments per animal, Fig. 8). After this, the number of segments in all individuals were counted once every two weeks over the course of six weeks. We found that the animals that were on the spinach regimen grew new segments at a notably faster rate (6 segments per week) than animals that were on the spirulina regimen (3.6 segments per week) (Fig.8).

**Figure 8.**
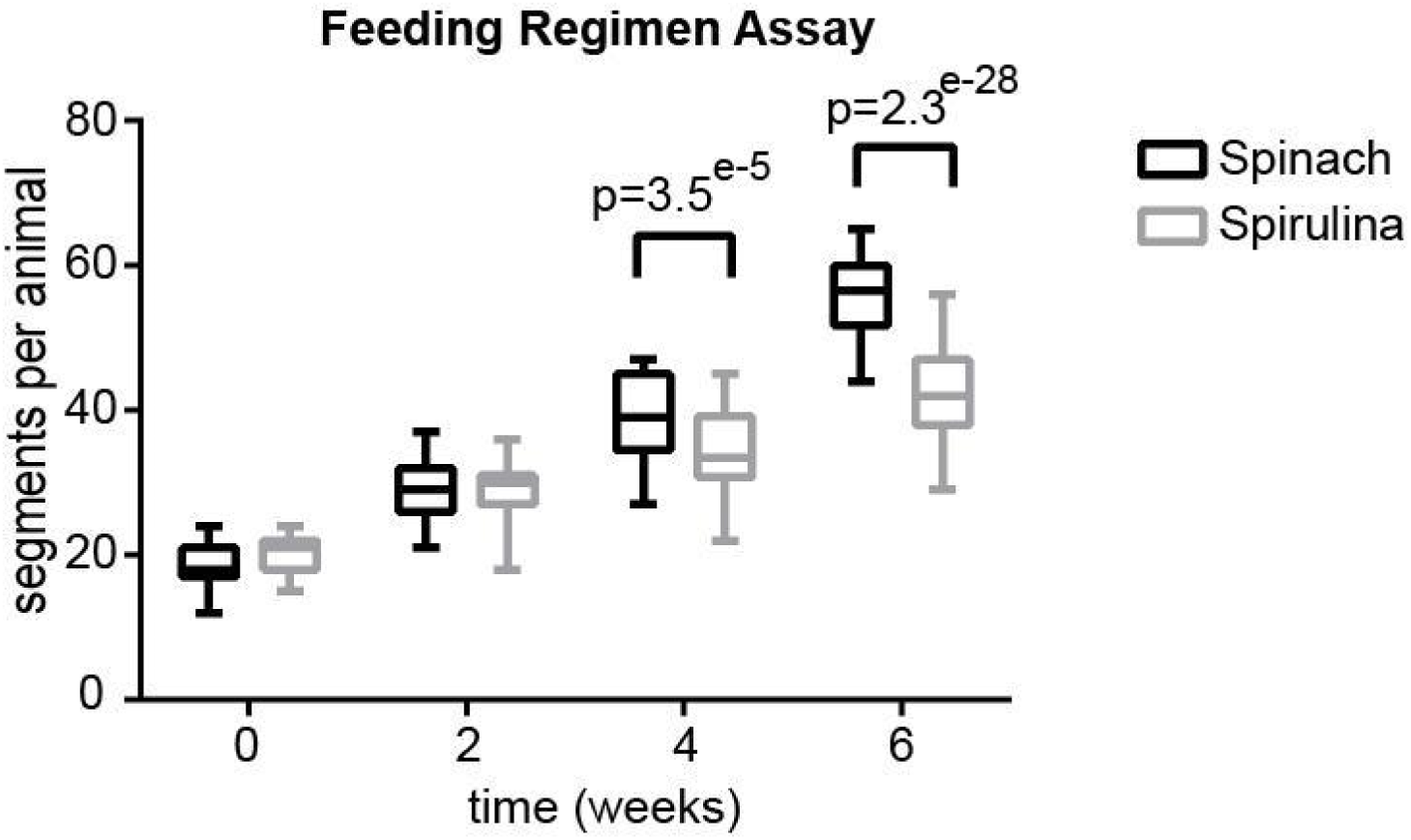
Feeding regimen comparison. Box-plot shows comparison of segment numbers in spirulina-fed and spinach-fed animals over the course of 6 weeks. To count the segments, each pair of parapodia was considered one segment, excluding the two pairs on the animal’s heads. On the posterior end of the worms, only segments carrying parapodia with visible chaetae were counted. Due to natural inter-individual variation in size of the animals, the average number of segments per animal differed slightly already at t=0, with an average of 19,2 segments per animal (SD=2,9). Over the course of the experiment, the spinach-fed animals showed a notably faster growth rate, adding an average of 6 segments per week per animal, compared to an average of 3.6 segments per week per animal in the spirulina-fed condition. A statistically significant difference in segment numbers was observed after 4 weeks (t-test, p=3.5e^−5^) and 6 weeks (t-test, p=2.3e^−28^) between animals that were under spinach vs spirulina regimens.

It is worth noting that the spinach-fed animals did not finish eating all the spinach provided each week and were therefore fed *ad libitum*, as opposed to the spirulina-fed animals that seemed to consume the algae provided in a relatively short time. This was observed by checking the color of the water (as in Suppl. Fig. 3A), which turned green upon feeding and cleared up again in the first 1-2 days after feeding. This indicates that the amount of food provided to these animals was not sufficient to grant optimal growth, which may explain the slower growth rate. In a future experiment varying amounts of spirulina cocktail will be tested and compared to the spinach regimen. In the culture boxes at the MBL (Woods Hole), we have extensive algal growth over the course of only a few weeks with no adverse effects (see Fig. 1G as an example). We suspect this algal film may act as an extra source of food for juvenile worms in addition to the spirulina cocktail, while during the above experiment (performed in Vienna cultures) the culture boxes did not develop such dense algal growth, which may have also affected the rate of growth. Overall, we find that the standardized spirulina-sera micron feeding regimen provides an easily-scalable and more accurately measured feeding method, which eventually yields health mature worms (see below).

### Feeding the young larvae with spirulina

After establishing that a defined spirulina diet can be used to maintain a culture of juvenile to mature worms, we next wondered whether a similar strategy could already be applied at earlier stages. Throughout early development, *P. dumerilii* larvae depend on yolk and four large lipid droplets, one contained in each macromere as their initial source of food [17]. These droplets are largely expended by the 3-segmented juvenile stage (around 5-7 dpf), and the developing worms must seek other energy sources in order to survive. Typically, in lab cultures feeding the larvae with live algae starts around 7-10 dpf. To our knowledge, several different algae species have been used successfully by different labs. Among these are *Tetraselmis marina, Tetraselmis sp*. (SAG no 3.98 from Department Experimental Phycology and Culture Collection of Algae (EPSAG)), *Isochrysis galbana* (also called *T-iso*), *and Tetraselmis chuii*. Ernest E. Just procured algae for his larval *P. dumerilii* cultures by scraping the bottoms of mariculture water tables for a “felt-like growth of diatoms and protozoa” [23].

Despite being a good nutritional source, live algae cultures are prone to contamination by protozoa or rotifers, and some of these organisms can end up over-populating *P. dumerilii* larvae cultures, causing poor culturing quality. In addition, keeping live algae cultures is time-consuming. We therefore tested if a spirulina-only cocktail (powdered spirulina in NFSW, 1.0 g/L) would provide an adequate substitute for the established algae cultures for feeding larvae.

Larvae from a single batch were split by volume into 3 groups around 24 hpf. At 7 dpf, they were transferred to culture boxes with 250 mL NFSW. Each group was fed with either: i) 10 mL spirulina-only ii) 25 mL spirulina-only iii) algae-only (10 mL live algae per box), 3 times per week (Mo, Wed, Fri). At 24 days, the animals were checked under the microscope for growth and development. Larvae in both spirulina-only boxes appeared healthy and, compared to the group fed with algae only, the spirulina-fed larvae appeared larger. The algae-fed culture box appeared very clean at this time, suggesting the amount of algae used may not have been enough (completely consumed). These cultures were not followed for the long-term effects of the feeding regime on maturation. However, the observation suggests it may be possible to grow larvae without requiring live algae cultures, while further and long-term testing of the spirulina-only regimen is still needed.

### Worm maturation under new culturing conditions

To ensure the proper functioning of our setup, we recorded the number of males, females, and worms of unknown gender over a seven-month period (Fig. 9). We did not explicitly look for mature animals outside of the scheduled maturation window. However, off-cycle maturations were recorded if we noticed mature animals while performing other husbandry tasks. Each cycle yielded an average of 53 mature worms (Fig. 9D), though the number of mature animals we found varied according to the number of worms which were in low density cultures (Suppl. Fig. 4). Our cultures began producing mature animals approximately five months after we received our initial *P. dumerilii* larvae (however, this corresponds to four months after light cycles started). The initial cultures were not introduced to the lighting regimen until approximately one-month post-fertilization (December 29, 2017) when the first low density boxes were made and transferred onto the shelving unit. It should therefore be noted that, typically, it is possible for mature animals to be procured more quickly (3-4 months post fertilization). At this time we had approximately 840 worms in low density cultures and found 22 mature worms. As we increased the number of low-density boxes in the shelving unit, the number of mature worms we found in the following months increased as expected. Three months after finding the first mature worms, we expanded our low-density cultures to house approximately 2000 worms, and around 60 mature worms were found (Fig. 9D, Suppl. Fig. 4).

**Figure 9.**
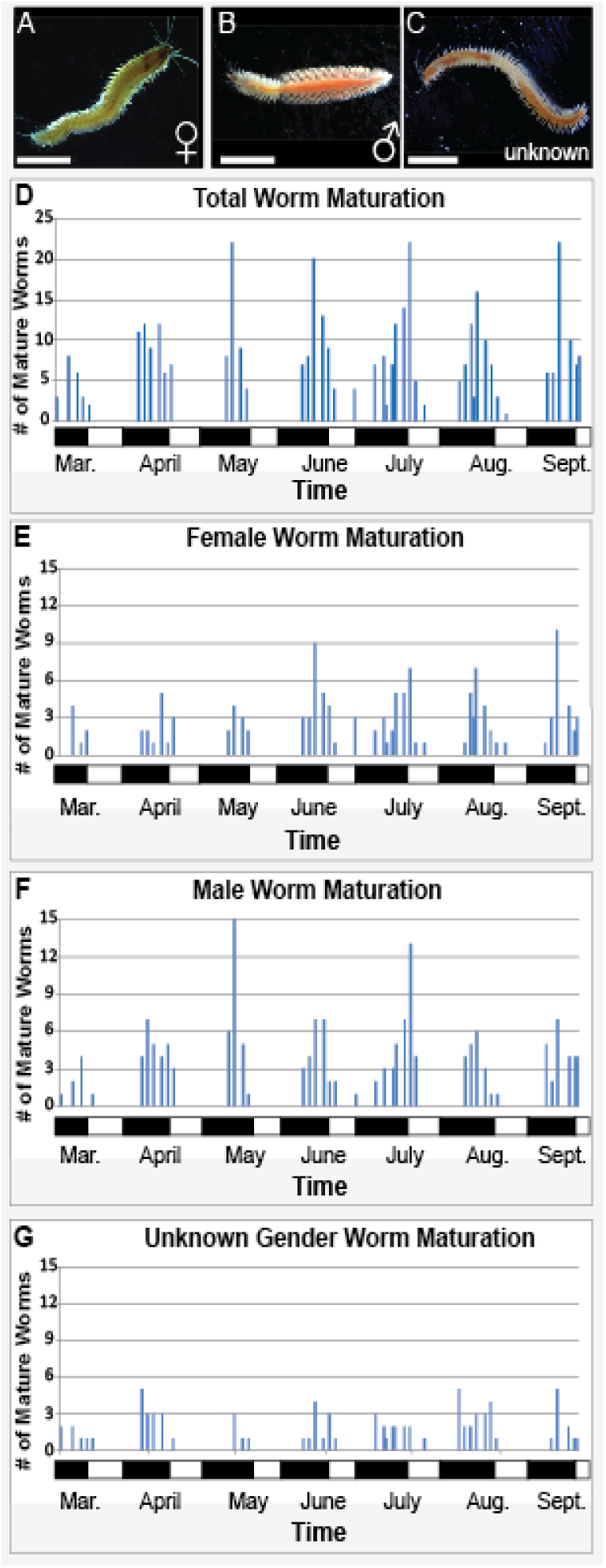
*P. dumerilii* Maturation. A) Mature male *P. dumerilii*. B) Mature female *P. dumerilii*. C) Mature *P. dumerilii* of unknown gender. D) Total number of mature *P. dumerilii* found for each given cycle. Culture boxes were checked 3 times per week during the maturation period. Black and white bars represent moon off and on cycles respectively (D-G). E) Number of female worms found during each cycle. F) Number of male worms found during each cycle. G) Number of worms of unknown gender found during each cycle. Note that cultures were systematically checked for mature worms only during the weeks maturation happens at a higher rate (for 2 weeks). We did not systematically check for mature worms outside of these 2 weeks. However, mature worms we came across outside of maturation weeks were noted as “off cycle” and are shown in the graphs. Bars on the X axis indicate Moon ON (white) and OFF (black) periods. Scale bars: 4 mm

Our cultures tended to produce a slightly greater number of mature males (average of 24/cycle) than females (average of 18/cycle) (Fig. 9E, F). We are uncertain if this is a product of our culturing conditions or if this mimics a more natural sex ratio for this species. Just observed the opposite in his laboratory cultures of *P. megalops*, where females outnumbered males, meanwhile he noted that in nature the reverse was the case (males outnumbered females) [23,28]. In addition, the peak maturation in our Woods Hole cultures starting at day 10 after the moon light is off differs slightly from what has been observed in some *P. dumerilii* strains in Vienna [16] and from the earliest reports from Ranzi on the Naples population [31,32]. This could be due to the method of scoring worms as mature: for example, here we used color change after metamorphosis and scored these worms as mature, while another method of scoring is by counting only the worms that have spawned as mature. Other possible reasons for the difference could be the light conditions (availability or lack of particular wavelengths), or genetics.

### Normal development under the reported culturing conditions

Under the culturing conditions we have reported here, we have been able to raise *P. dumerilii* for several generations at the MBL, and obtained normally-developing embryos, larvae and juveniles (Fig. 10). We have also injected fertilized eggs with mRNA and successfully reproduced the results obtained with past *P. dumerilii* cultures [5] (Fig. 10F, F’). Different from other labs working with *P. dumerilii*, we have observed jelly production by fertilized oocytes to be longer (until around 1,5 hpf compared with the cultures in Paris and Vienna which typically release jelly until around 1 hpf). We suspect the slightly lower salinity levels of the sea water in Woods Hole, or some undetermined factors in the sea water may be the cause of the jelly production to last longer than usual in our cultures. We have also observed juvenile worms to regenerate successfully at a similar rate to the published stages of regeneration for *P. dumerilii* (results not shown) [6].

**Figure 10.**
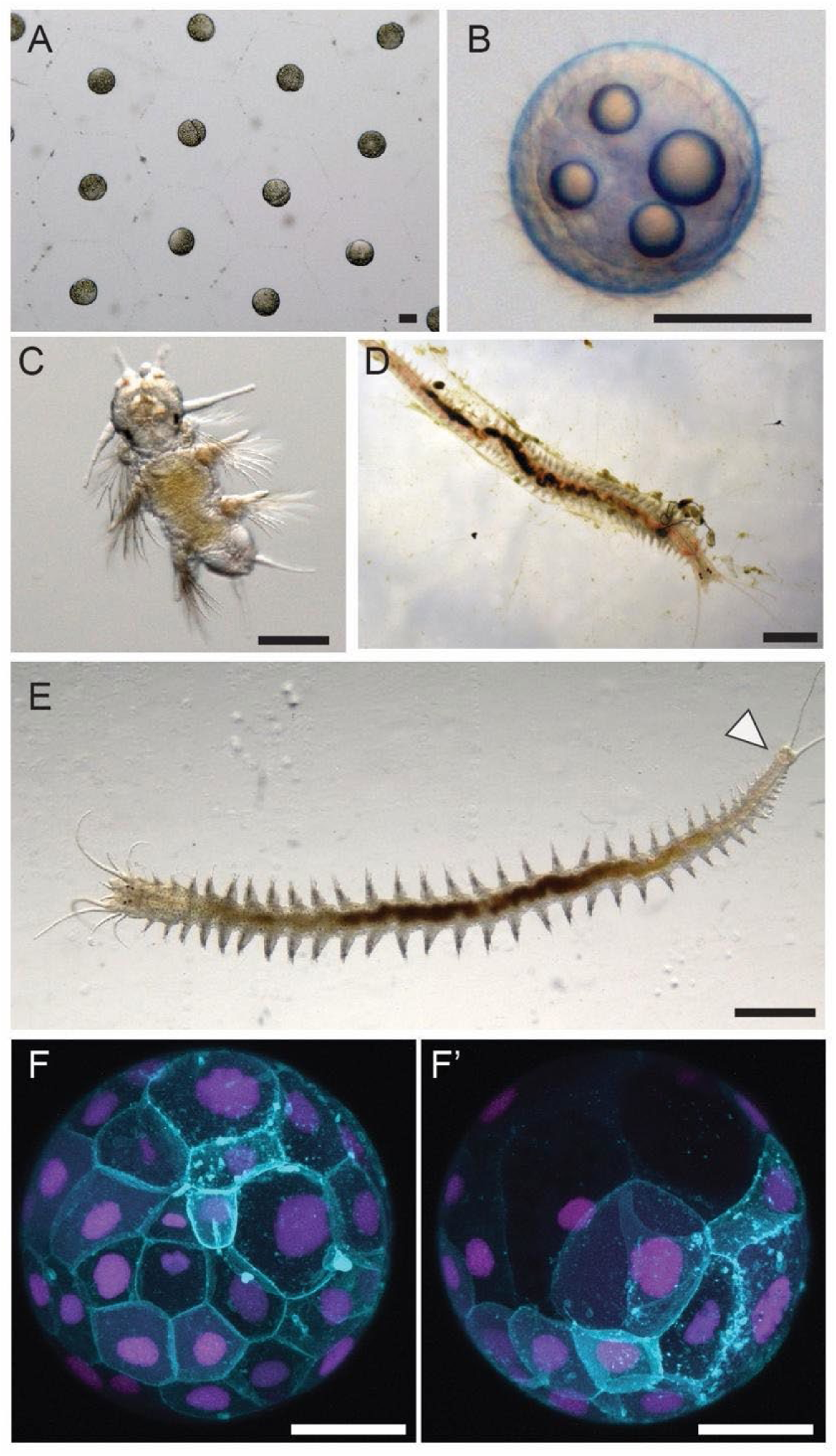
.dumerilii development. A) *P. dumerilii* embryos at 2 hpf (hours post fertilization). Following fertilization, embryos secrete a protective jelly which appears as a hexagonal matrix. B) At 17 hpf *P. dumerilii* larvae reach the the prototroch stage. The larvae develop a band of cilia by 12 hpf which allows them to start swimming. C) *P. dumerilii* larva at 9 days post fertilization. D) Adults burrow in tubes made from a matrix of secreted mucus (Daly, 1973). E) *P. dumerilii* adults continue adding body segments throughout their lives from a posterior growth zone (arrowhead). F-F’) Embryos injected with EGFP-caax and Histone-mCherry mRNA were imaged around 7 hpf using a confocal scope. Scale bars: 100 µm (A-C), 5 mm (D,E), 50 µm (F-F’).

### Future directions

Future improvements to culturing *Platynereis* and other aquatic organisms remain. Integrating temperature control to the shelving setup itself (for example, by using a cooling device on each shelf) instead of controlling the temperature of the culture room will allow greater flexibility and reliability. Determining why natural sea water is needed for culturing *Platynereis* embryos and larvae, and identifying the organic and inorganic factors that make the natural sea water critical for optimal culturing outcomes may allow developing a working artificial seawater recipe and switching to artificial sea water entirely. Finally, determining whether there are more specific light spectral needs to improve *Platynereis* maturation rates will be helpful to have faster maturation times.

The benefits of the system we developed are not only scalability, but also the possibility to control food input more accurately, and the possibility to vary environmental parameters more flexibly. These are crucial components of culturing especially, as the living systems and their biology is directly affected by nutrition, and often by light cycles. Being able to control these parameters and standardize them will lead to easier reproducibility of experiments and techniques by different labs. We also envision that this culture setup design, detailed blueprints, and standardized feeding methods presented here will be beneficial to not only the community of labs that use *P. dumerilii* as a research organism, but also those labs that are interested in having *P. dumerilii* at a small scale for specific experiments and/or educational purposes, as well as for labs studying other aquatic invertebrates.

## Acknowledgements

We thank John Carr, Kate Dever, Lindsey DeMelo and the rest of the MRC staff at the MBL for their help in building the shelving setup and culture husbandry; we thank Jon Henry, and colleagues working on *P. dumerilii* for their helpful suggestions; and Guillaume Balavoine for providing the *P. dumerilii* larvae to start our cultures.

## Author Contributions

BDO and JG designed the shelving setup; EK, AWS and BDO collected the data; EK, AWS, FR, and BDO analyzed the data; EK, AWS, and BDO drafted the manuscript; FR and BDO edited the manuscript. All authors read and approved the manuscript.

## SUPPLEMENTARY FIGURES AND LEGENDS

**Supplementary Figure 1.**
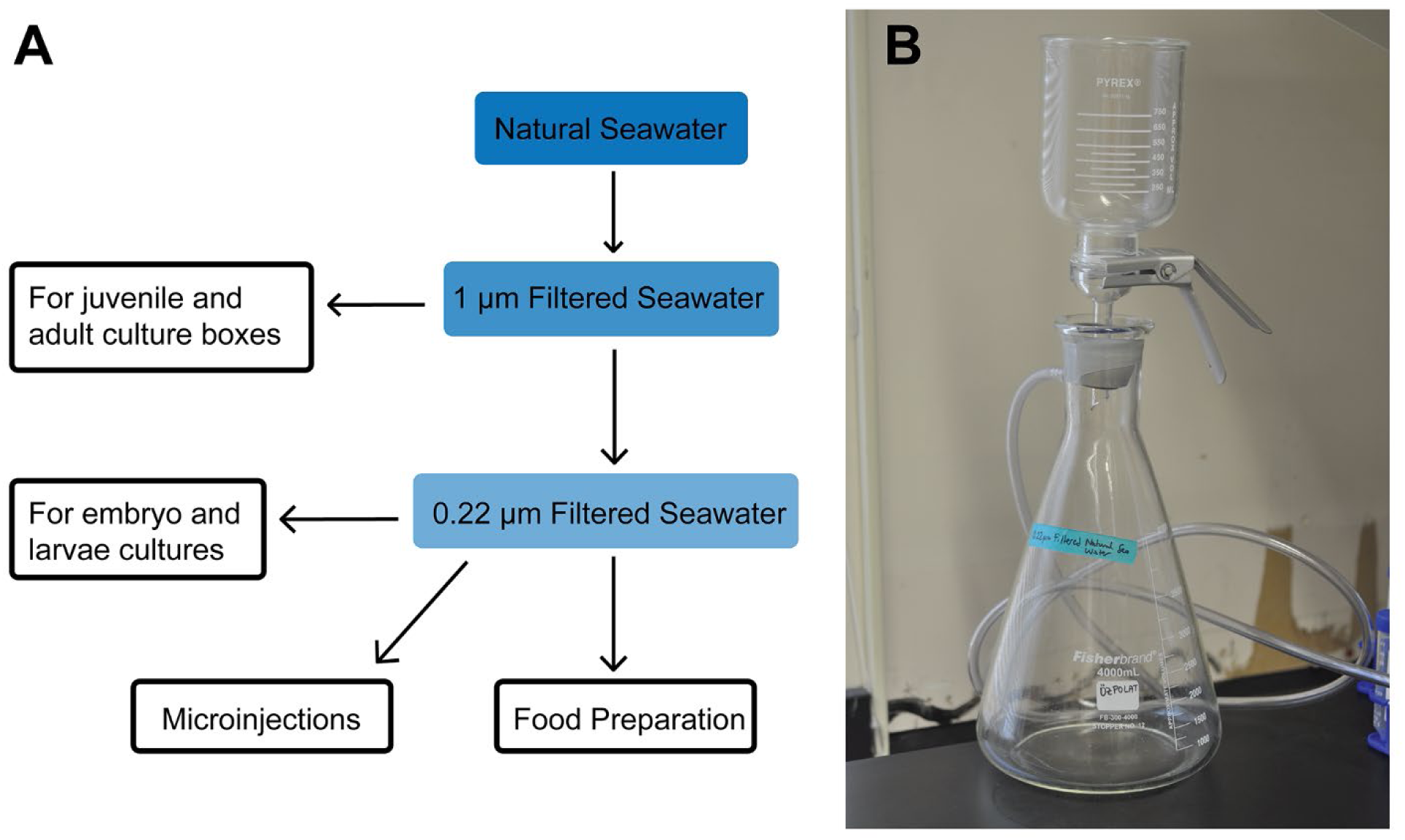
Water Filtration. A) Schematic of seawater filtering process and what each level of filtration is used for. B) Our 0.22 µm filtration set up. This system is more economical than plastic filter bottles. See supplementary spreadsheet for catalog numbers and ordering information.

**Supplementary Figure 2.**
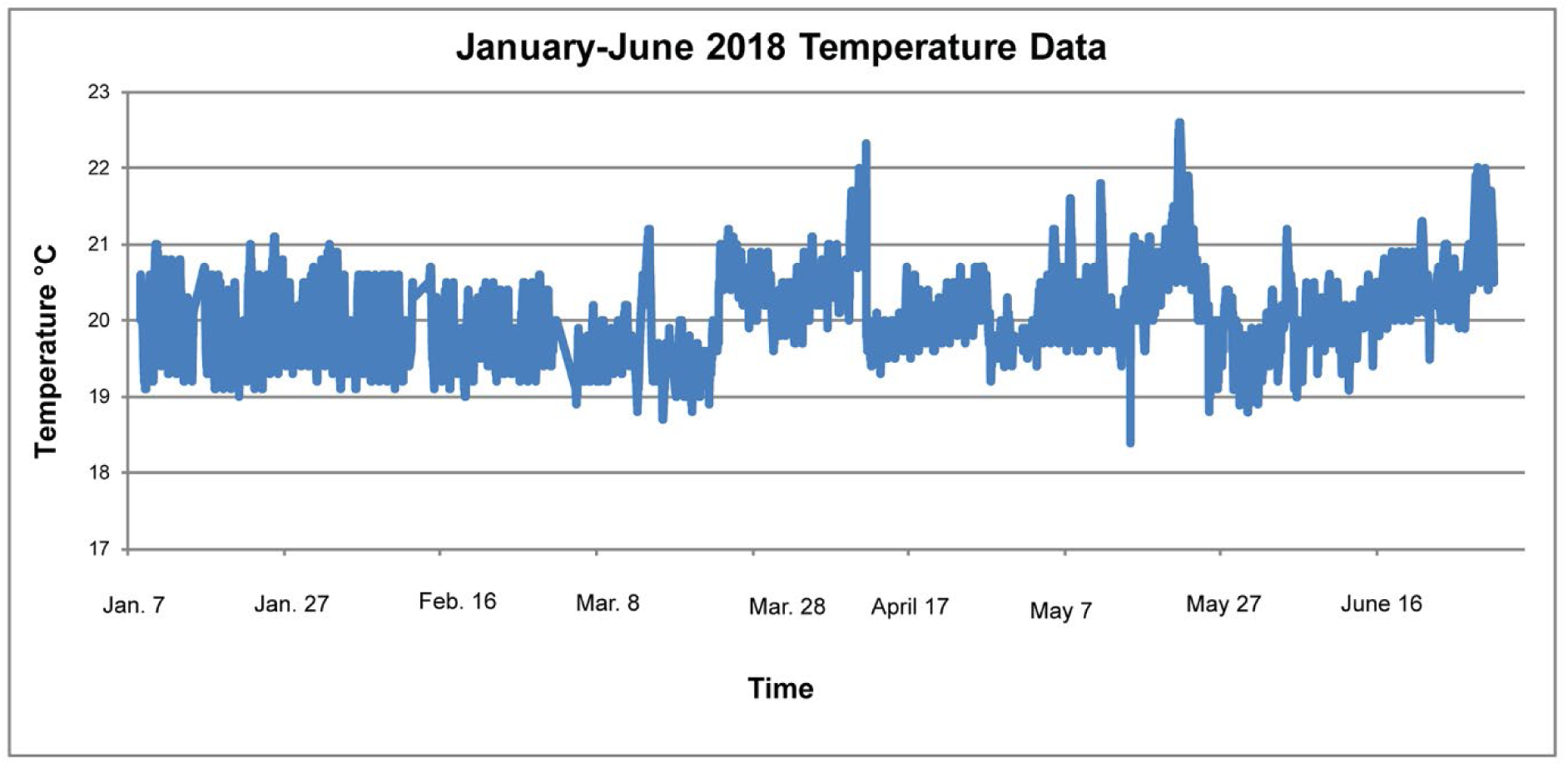
Temperature. Cultures were kept at 20°C with some variability in the spring and summer months due to fluctuations in the temperature of the building. A portable air conditioning unit was used to help better regulate temperature during the summer months after the peaking of temperature above 22°C in April and May.

**Supplementary Figure 3.**
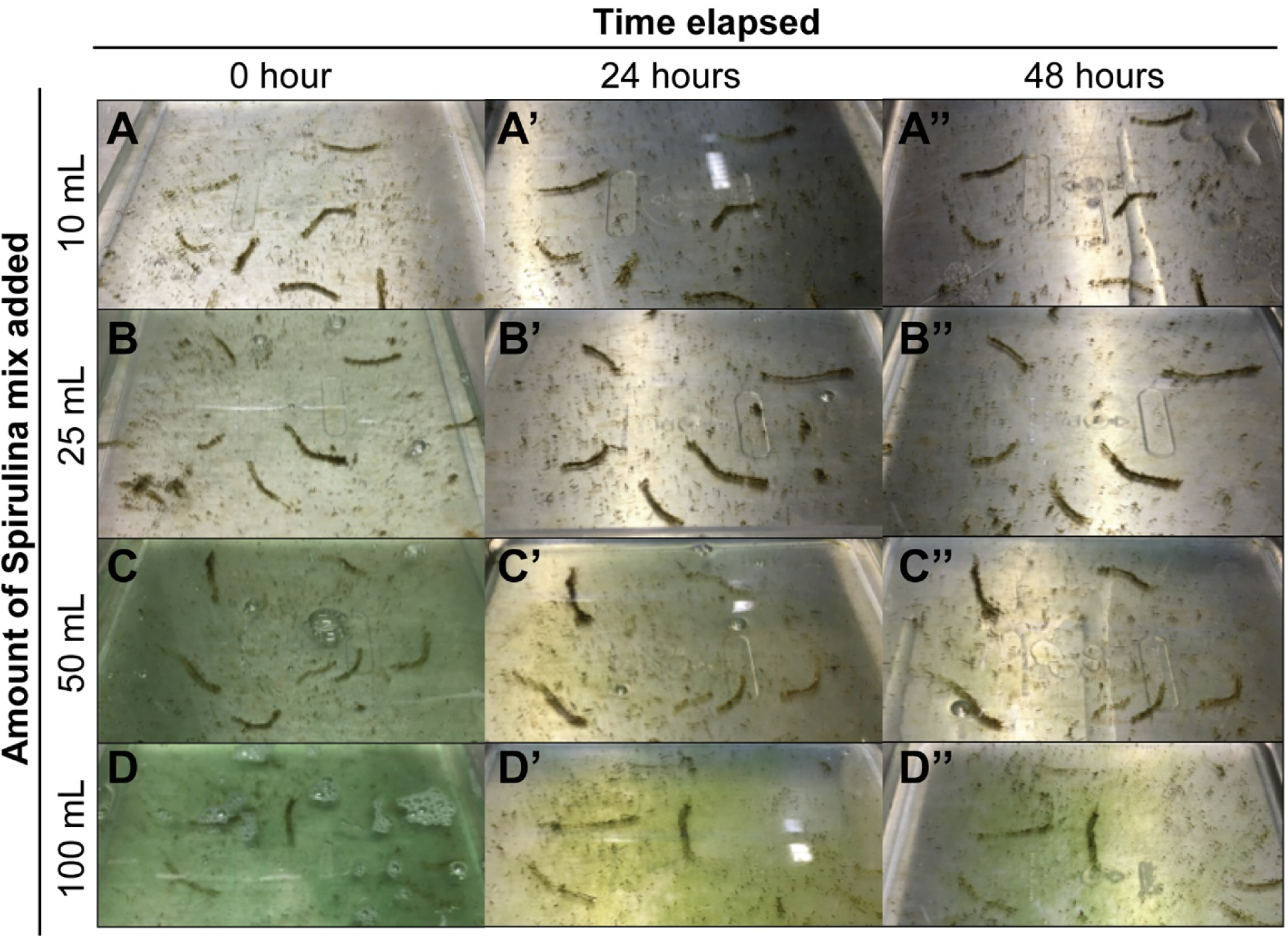
Spirulina gradient. To test which concentrations of spirulina cocktail can be fed safely to a box of 10 animals without rotting and spoiling the water, four boxes of 10 animals each were fed the following amounts of spirulina cocktail: 10, 25, 50 and 100 mL. Images were taken directly after feeding, 24 hours later and 48 hours later.

**Supplementary Figure 4.**
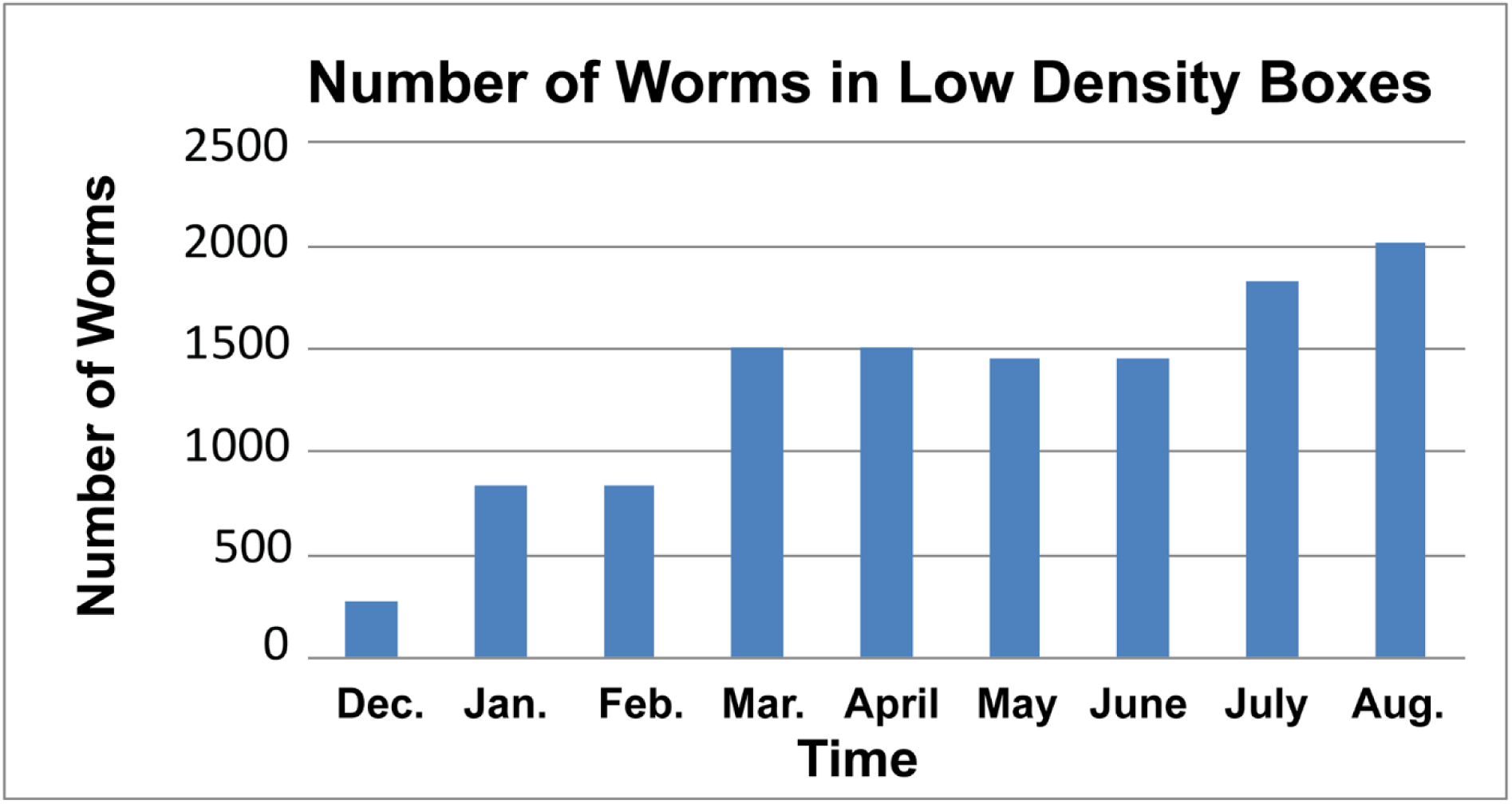
Number of worms in low density boxes. The graph shows the number of worms living in low density cultures (30 worms/small Sterilite box) (for the period December 2017-August 2018). The number of mature animals found each cycle increased (Fig. 9D) as more low density cultures were established. These numbers (along with mature animal numbers in Fig. 9) can be used as a guide to scale up or down low density culture boxes, for obtaining mature animal numbers desired.

## REFERENCES

1. Wilson EB. The cell-lineage of Nereis. A contribution to the cytogeny of the annelid body. J Morphol. Wiley Subscription Services, Inc., A Wiley Company; 1892;6: 361–480.

2. Fischer A, Dorresteijn A. The polychaete Platynereis dumerilii (Annelida): a laboratory animal with spiralian cleavage, lifelong segment proliferation and a mixed benthic/pelagic life cycle. Bioessays. 2004;26: 314–325.

3. Williams EA, Jékely G. Towards a systems-level understanding of development in the marine annelid Platynereis dumerilii. Curr Opin Genet Dev. 2016;39: 175–181.

4. Balavoine G. Segment formation in Annelids: patterns, processes and evolution. Int J Dev Biol. 2014;58: 469–483.

5. Özpolat BD, Handberg-Thorsager M, Vervoort M, Balavoine G. Cell lineage and cell cycling analyses of the 4d micromere using live imaging in the marine annelid Platynereis dumerilii. Elife. 2017;6. doi:10.7554/eLife.30463

6. Planques A, Malem J, Parapar J, Vervoort M, Gazave E. Morphological, cellular and molecular characterization of posterior regeneration in the marine annelid Platynereis dumerilii. Dev Biol. 2018; doi:10.1016/j.ydbio.2018.11.004

7. Gazave E, Béhague J, Laplane L, Guillou A, Préau L, Demilly A, et al. Posterior elongation in the annelid Platynereis dumerilii involves stem cells molecularly related to primordial germ cells. Dev Biol. 2013;382: 246–267.

8. Ayers T, Tsukamoto H, Gühmann M, Veedin Rajan VB, Tessmar-Raible K. A Go-type opsin mediates the shadow reflex in the annelid Platynereis dumerilii. BMC Biol. 2018;16: 41.

9. Schenk S, Bannister SC, Sedlazeck FJ, Anrather D, Minh BQ, Bileck A, et al. Combined transcriptome and proteome profiling reveals specific molecular brain signatures for sex, maturation and circalunar clock phase. Elife. 2019;8. doi:10.7554/eLife.41556

10. Lauri A, Brunet T, Handberg-Thorsager M, Fischera. HL, Simakov O, Steinmetz PRH, et al. Development of the annelid axochord: Insights into notochord evolution. Science. 2014; doi:10.1126/science.1253396

11. Brunet T, Fischer AHL, Steinmetz PRH, Lauri A, Bertucci P, Arendt D. The evolutionary origin of bilaterian smooth and striated myocytes. Elife. eLife Sciences Publications Limited; 2016;5: e19607.

12. Ackermann C, Dorresteijn A, Fischer A. Clonal domains in postlarval Platynereis dumerilii (Annelida: Polychaeta). J Morphol. 2005;266: 258–280.

13. Nakama AB, Chou H-C, Schneider SQ. The asymmetric cell division machinery in the spiral-cleaving egg and embryo of the marine annelid Platynereis dumerilii. BMC Dev Biol. 2017;17: 16.

14. Rebscher N, Lidke AK, Ackermann CF. Hidden in the crowd: primordial germ cells and somatic stem cells in the mesodermal posterior growth zone of the polychaete Platynereis dumerillii are two distinct cell populations. Evodevo. 2012;3: 9.

15. Vergara HM, Bertucci PY, Hantz P, Tosches MA, Achim K, Vopalensky P, et al. Whole-organism cellular gene-expression atlas reveals conserved cell types in the ventral nerve cord of Platynereis dumerilii. Proc Natl Acad Sci U S A. 2017;114: 5878–5885.

16. Zantke J, Ishikawa-Fujiwara T, Arboleda E, Lohs C, Schipany K, Hallay N, et al. Circadian and Circalunar Clock Interactions in a Marine Annelid. Cell Rep. The Authors; 2013;5: 99–113.

17. Fischer AH, Henrich T, Arendt D. The normal development of Platynereis dumerilii (Nereididae, Annelida). Front Zool. 2010;7: 31.

18. Backfisch B, Veedin Rajan VB, Fischer RM, Lohs C, Arboleda E, Tessmar-Raible K, et al. Stable transgenesis in the marine annelid Platynereis dumerilii sheds new light on photoreceptor evolution. Proc Natl Acad Sci U S A. 2013;110: 193–198.

19. Zantke J, Bannister S, Rajan VBV, Raible F, Tessmar-Raible K. Genetic and genomic tools for the marine annelid Platynereis dumerilii. Genetics. 2014;197: 19–31.

20. Veedin-Rajan VB, Fischer RM, Raible F, Tessmar-Raible K. Conditional and specific cell ablation in the marine annelid Platynereis dumerilii. PLoS One. 2013;8: e75811.

21. Bannister S, Antonova O, Polo A, Lohs C, Hallay N, Valinciute A, et al. TALENs mediate efficient and heritable mutation of endogenous genes in the marine annelid Platynereis dumerilii. Genetics. 2014;197: 77–89.

22. Achim K, Eling N, Vergara HM, Bertucci PY, Brunet T, Collier P, et al. Whole-body single-cell sequencing of the Platynereis larva reveals a subdivision into apical versus non-apical tissues [Internet]. bioRxiv. 2017. p. 167742. doi:10.1101/167742

23. Just EE. On Rearing Sexually Mature Platynereis megalops from Eggs. Am Nat. [University of Chicago Press, American Society of Naturalists]; 1922;56: 471–478.

24. Just EE. The Relation of the First Cleavage Plane to the Entrance Point of the Sperm. Biol Bull. Marine Biological Laboratory; 1912;22: 239–252.

25. Hempelmann F. Zur Naturgeschichte von Nereis dumerilii. Aud et Edw Zoologica. 1911;25: 1–135.

26. Fischer A, Dorresteijn A. Culturing Platynereis dumerilii [Internet]. Available: http://www.staff.uni-giessen.de/~gf1307/breeding.htm

27. García-Alonso J, Smith BD, Rainbow PS. A compacted culture system for a marine model polychaete (Platynereis dumerilii). Pan-American Journal of Aquatic Sciences. 2013;8: 142–146.

28. Just EE. Breeding Habits of the Heteronereis Form of Platynereis megalops at Woods Hole, Mass. Biol Bull. Marine Biological Laboratory; 1914;27: 201–212.

29. Raible F, Takekata H, Tessmar-Raible K. An Overview of Monthly Rhythms and Clocks. Front Neurol. Frontiers Media SA; 2017;8: 189.

30. Cho E, Oh JH, Lee E, Do YR, Kim EY. Cycles of circadian illuminance are sufficient to entrain and maintain circadian locomotor rhythms in Drosophila. Sci Rep. 2016;6: 37784.

31. Ranzi S. Ricerche sulla biologia sessuale degli Anellidi. Pubbl Staz Zool Napoli. 1931;11: 271–292.

32. Ranzi S. Maturita sessuale degli Anellidi e fasi lunari. Boll Soc Ital Biol Sper. 1931;6: 18.

