## Supplementary File 2 for "A scalable culturing system for the marine annelid *Platynereis dumerilii*"

**Setting Up the Timer for Light Control**

Manufacturer’s manual can be found [here](http://www.nsiindustries.com/UserFiles/Documents/Product/MLI-199(B).DG180.DG280.103013.pdf) if needed.

To setup Sun (CH1) and Moon (CH2) cycles, follow this:

1) On the screen that says “SCH” press ENTER:
   a. Schedule 01: CH1 ON 06:00
   b. Schedule 02: CH1 OFF 22:00

This means the channel will be on at 6 AM and off at 10 PM

c. Schedule 03: CH2 ON 18:00
d. Schedule 04: CH2 OFF 06:00

This means the channel will be on at 6 PM and off at 6 AM

2) On the screen that says “SKIP” press D/ON:
   a. CH1 28:00 (SUN)

This means the channel will be ON 28 days during the hours set in the schedule above, and OFF 00 days.

b. Press CH Select
c. CH2 08:20 (MOON)

This means the channel will be ON during the first 8 days at the hours set in the schedule above (and it will be OFF during the remaining 20 days).

Cycle screen in the timer is not used for this setup.

The timer is the grey panel on the left in the picture below.

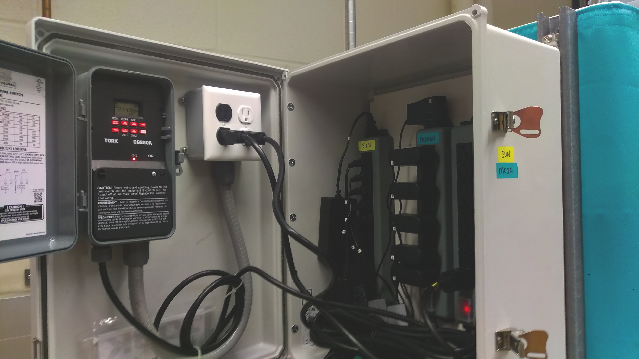

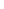

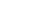

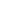

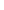

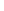

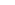

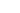

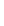

The first screen on the timer shows the current time and what channels are ON or OFF:

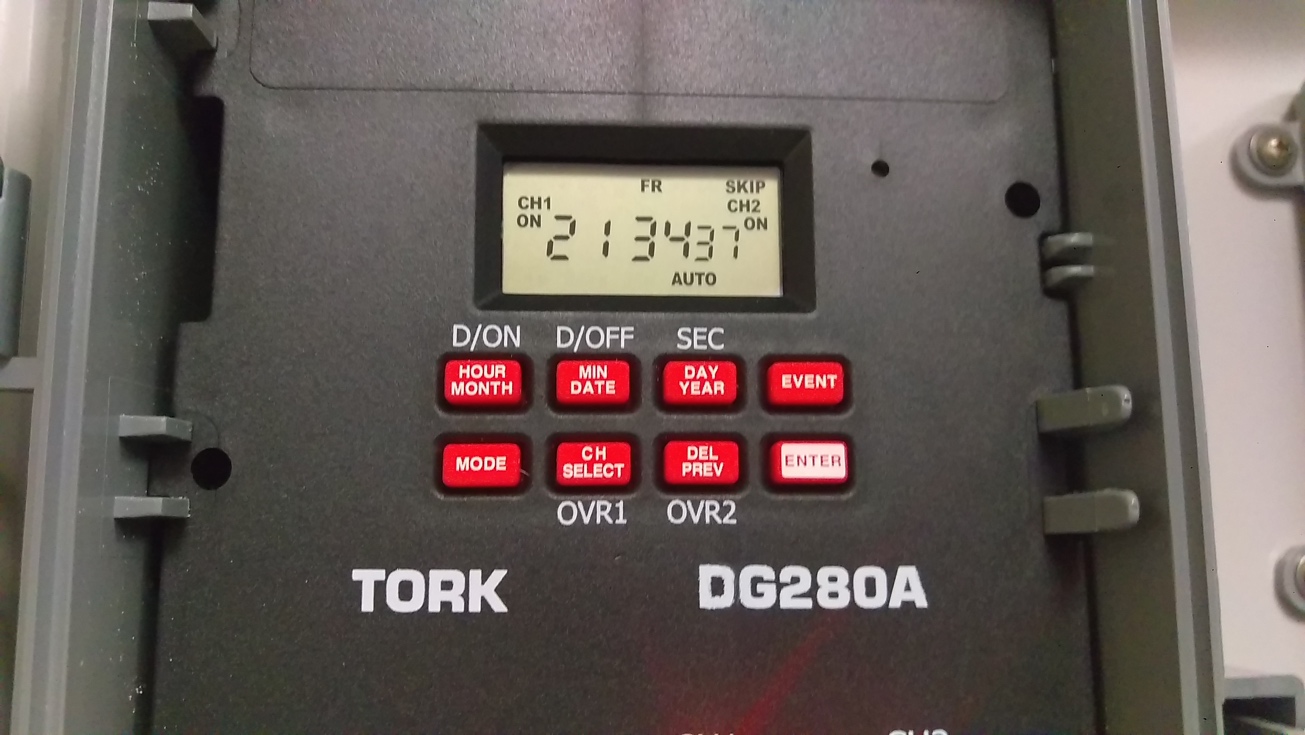

Press MODE to go through different setup options. When MODE is pressed, the next screen shows the current time:

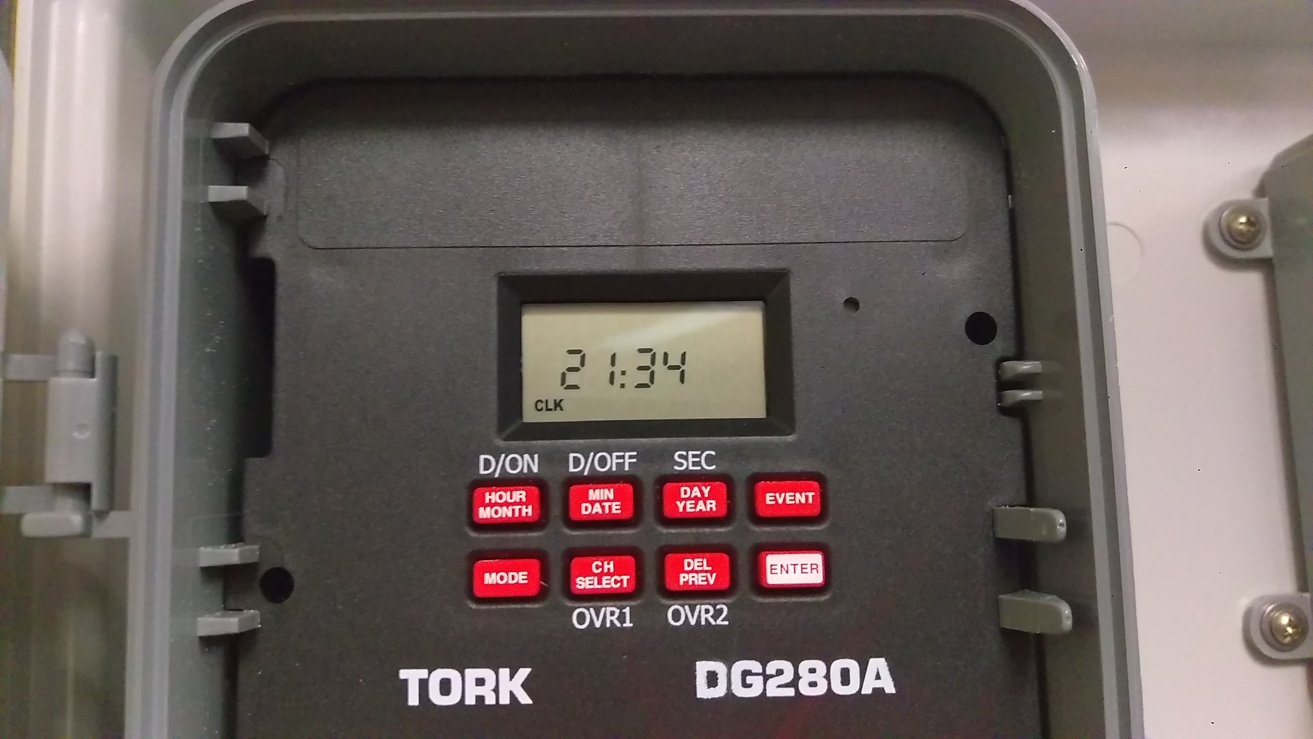

Next screen shows the current date (December 28^th^, 2018)

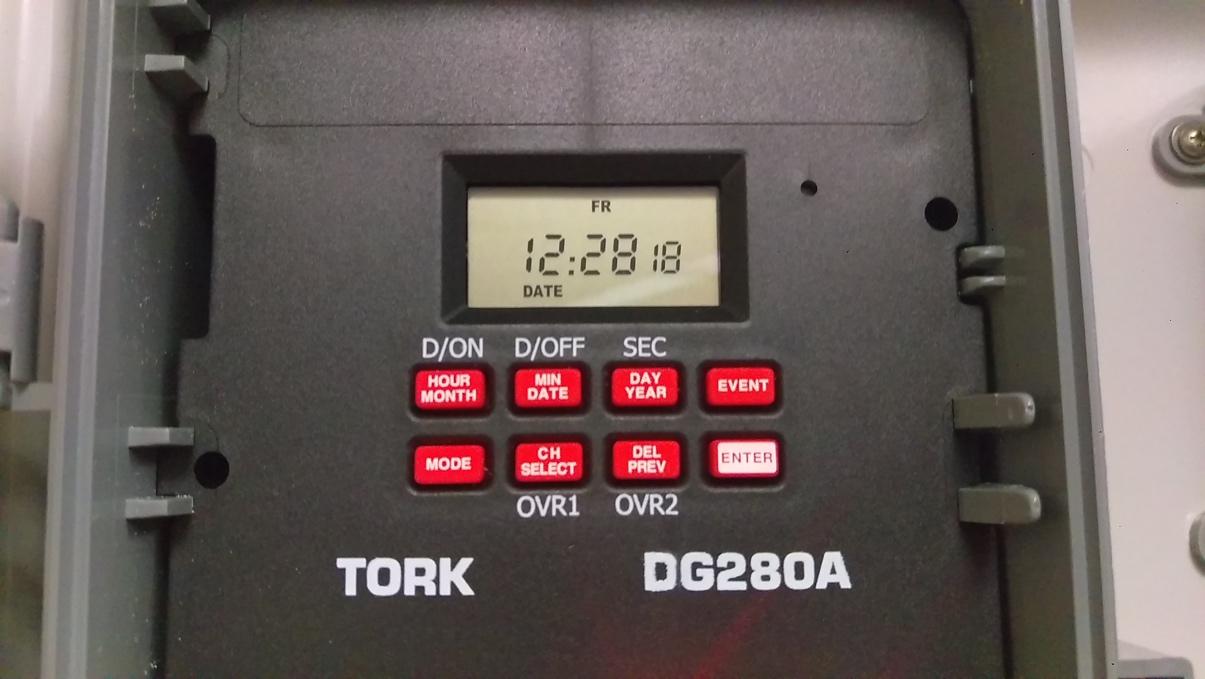

Screen for setting daylight saving time (not necessary to have it ON, we leave it OFF)

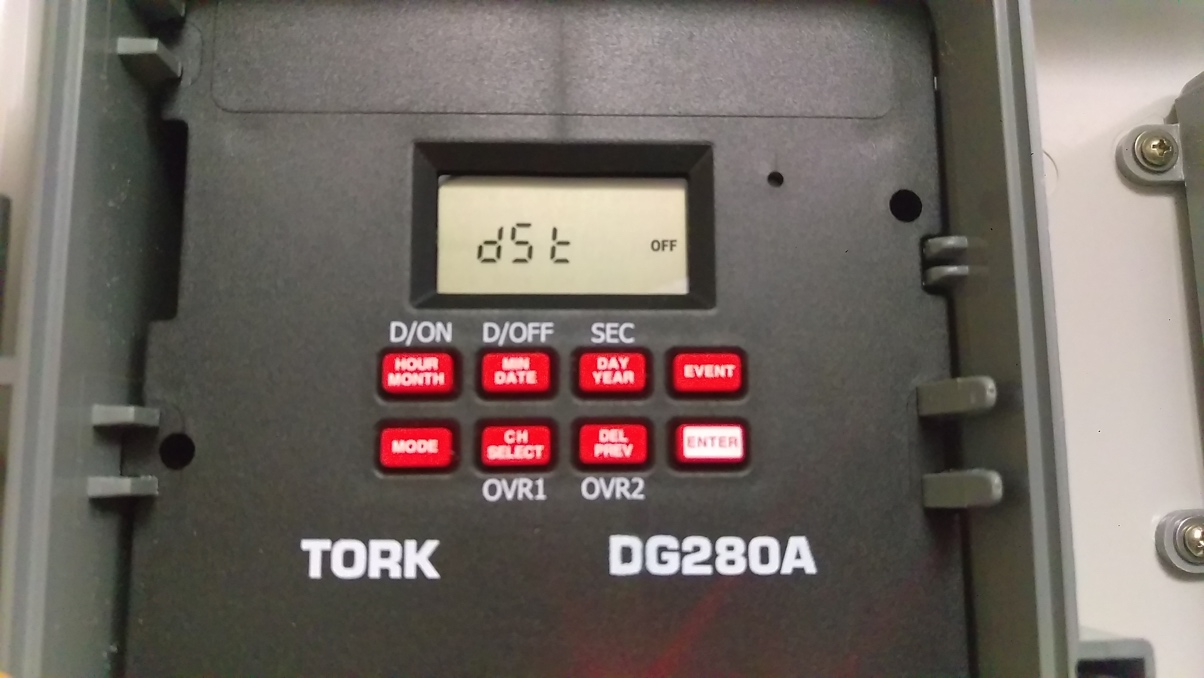

This is the CYCLE screen, no adjustment necessary for the *Platynereis* culture setup:

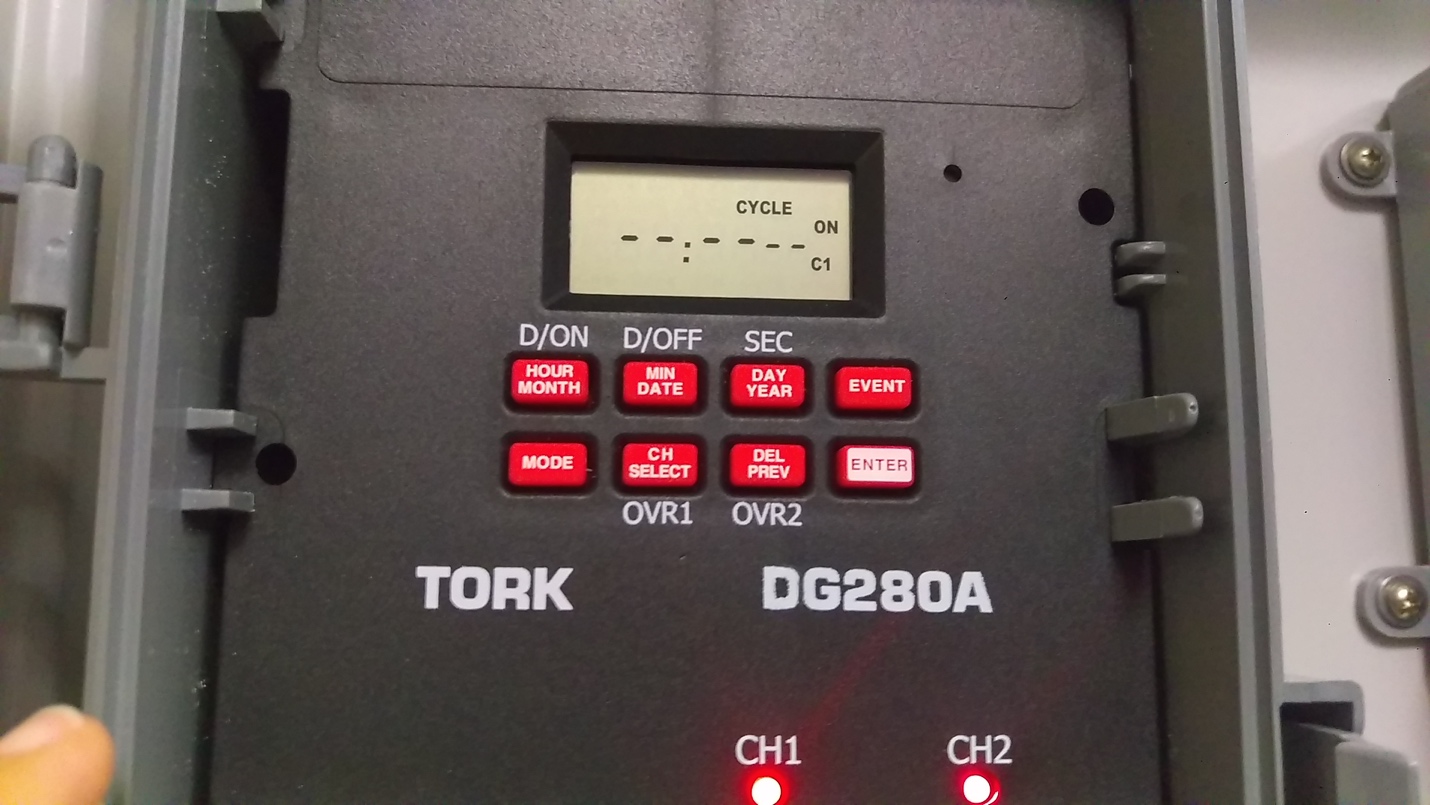

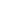

Next screen shows the SKIP settings:

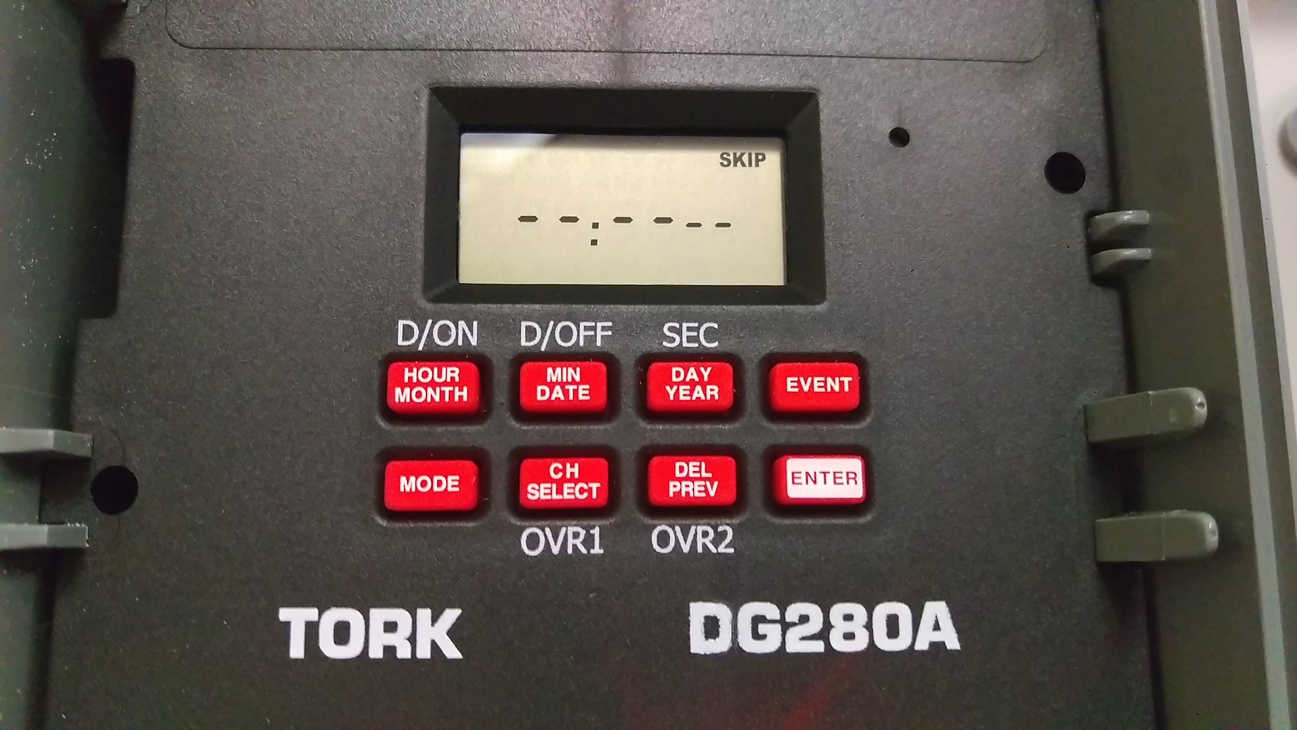

Press CH SELECT to see the current settings.

CH1 settings should appear as follows:

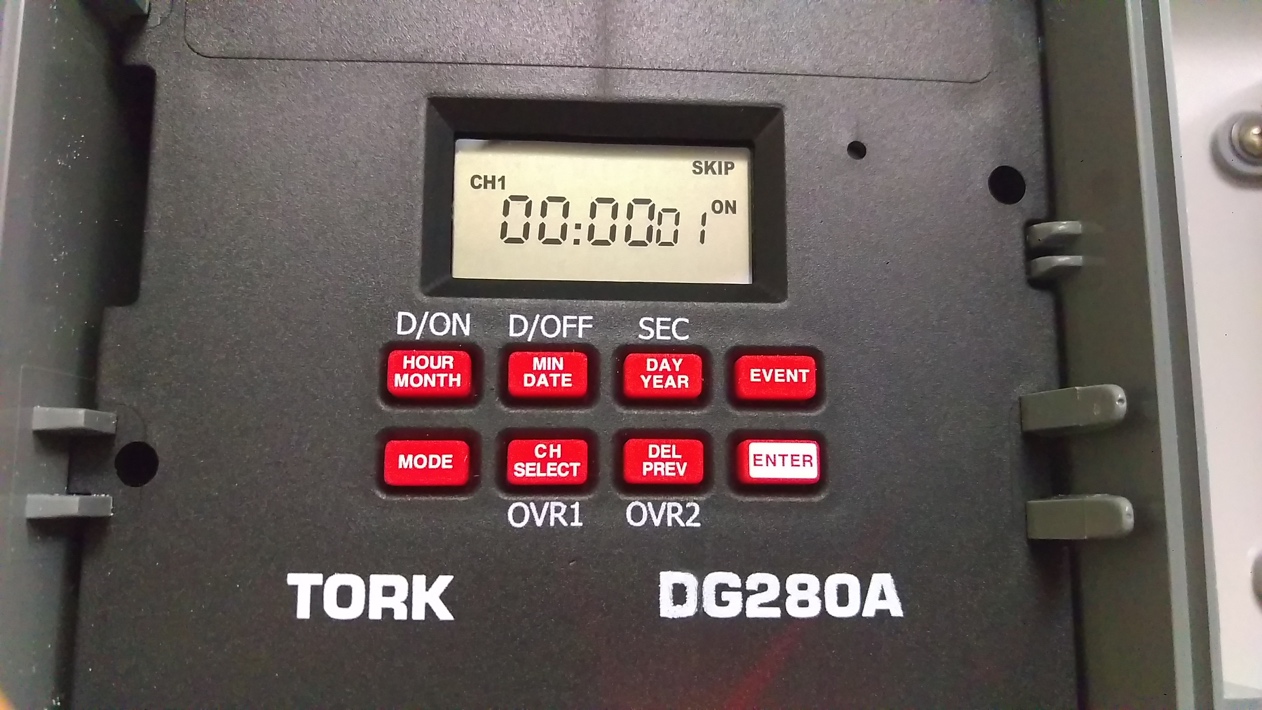

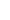

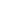

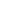

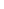

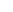

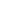

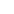

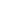

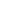

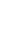

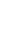

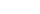

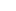

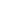

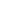

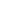

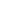

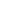

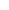

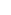

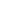

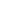

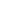

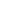

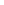

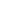

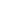

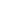

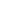

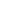

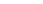

CH2 settings should appear as follows when CH SELECT is pressed again:

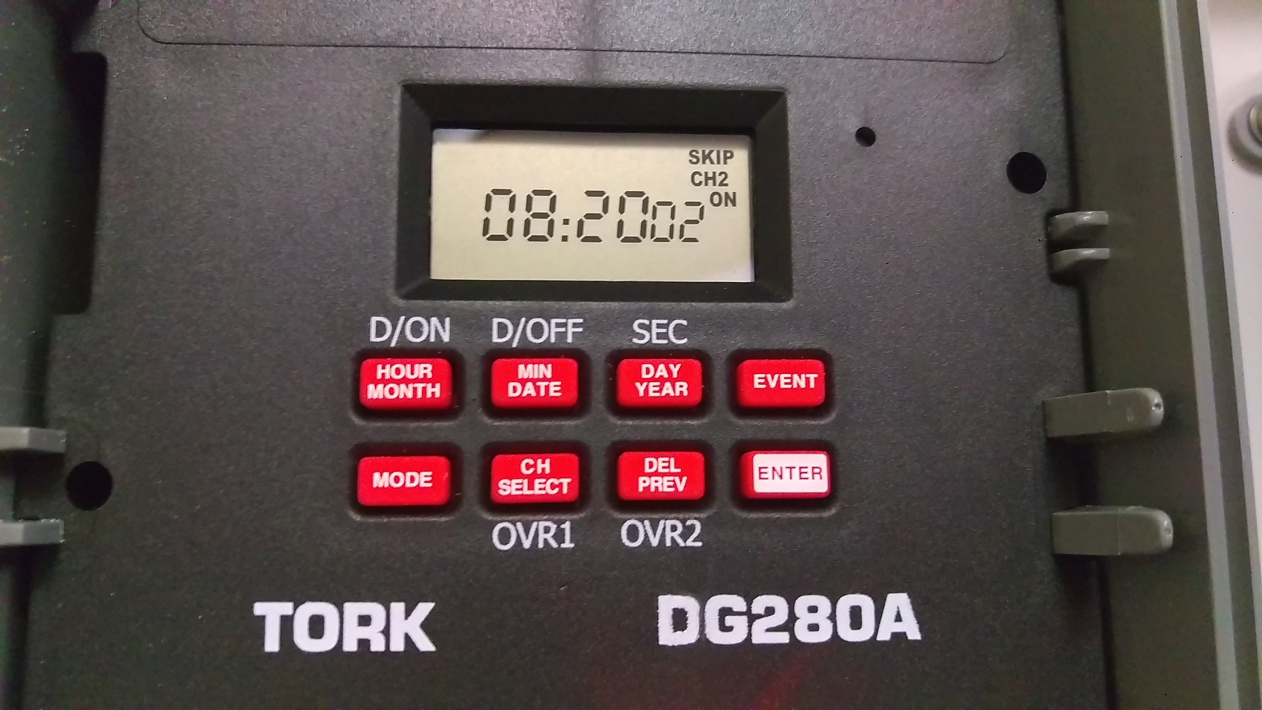

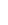

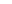

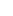

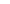

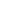

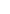

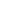

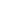

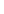

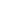

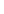

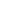

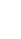

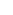

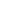

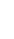

Next screen is the SCH (schedule) screen.

This (Schedule 01) shows ON schedule for CH1:

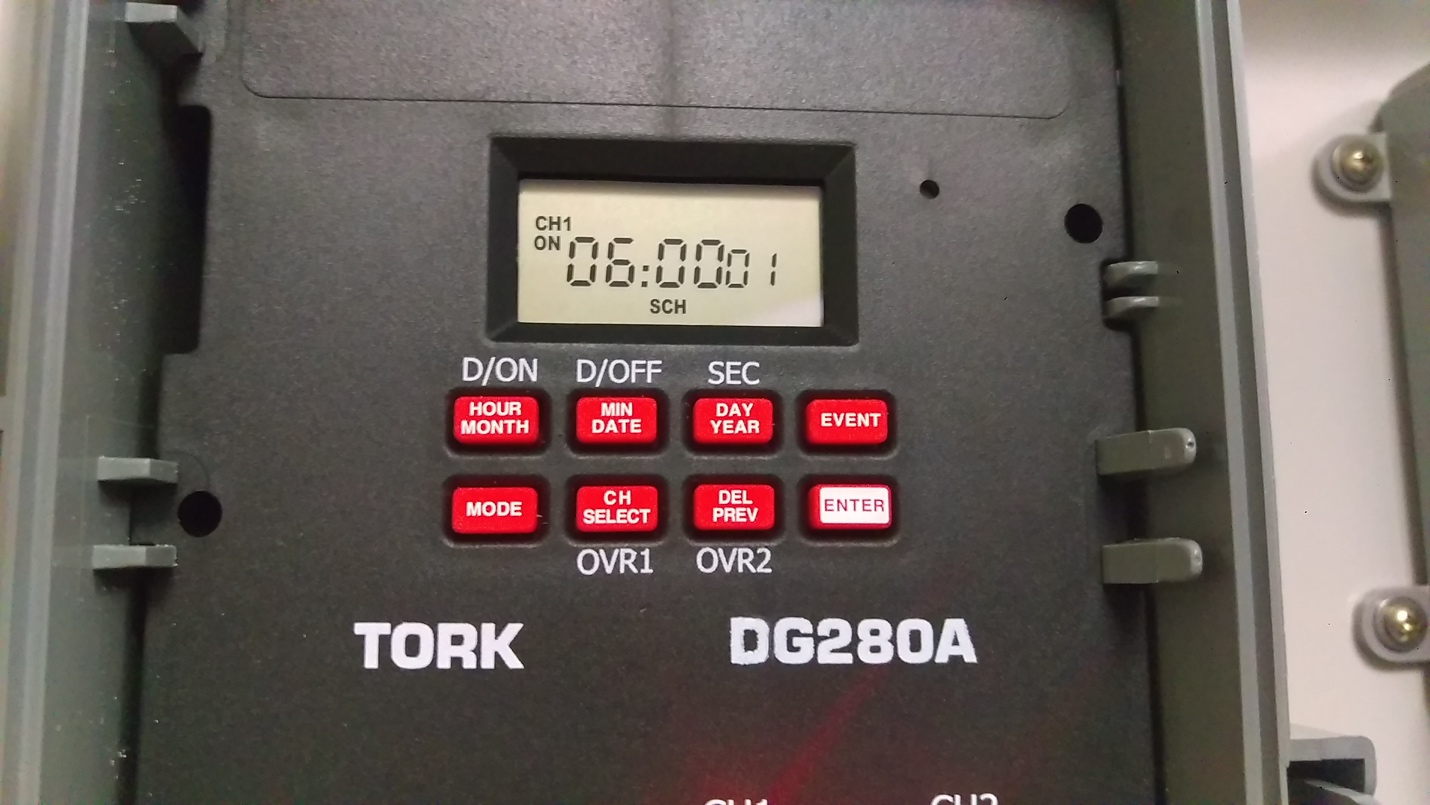

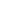

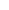

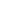

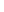

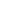

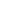

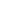

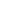

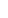

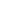

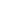

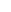

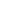

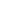

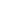

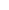

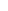

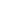

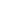

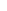

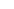

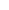

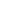

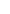

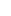

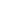

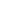

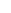

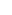

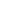

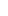

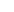

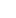

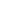

Press ENTER to see (and if needed, change) the rest of the schedule settings.

This (Schedule 02) shows the OFF schedule for CH1.

Press ENTER again. This (Schedule 03) shows the ON schedule for CH2.

Press ENTER again. This (Schedule 04) shows the OFF schedule for CH2.
